# Age-associated B cells (ABCs) develop from a CNS-localized progenitor pool into a pro-inflammatory phenotype after stroke

**DOI:** 10.64898/2026.03.02.709112

**Authors:** Annabel M. McAtee, Mathew Kenwood, Thomas Ujas, Mary K. Colson, Julie Watkins, Edric D. Winford, Katherine M. Cotter, Daimen Britsch, Dene Betz, Jadwiga Turchan-Cholewo, Jenny Lutshumba, Connor Stuart, Gargi Shah, Loganann Runice, Mark Ebbert, Salvatore Cherra, Peter T. Nelson, Jamie L. Sturgill, Nancy L. Monson, Mark P. Goldberg, Ann M. Stowe

## Abstract

Aging and age-related diseases like ischemic stroke induce chronic lymphocyte recruitment into the central nervous system (CNS). Conflicting effects on post-stroke functional recovery, however, are secondary to the differences in responding lymphocyte populations that shift immunophenotype with both ischemic injury and age. To better define CNS-localized B cell subsets, we used flow cytometry, single-cell RNA sequencing, and B cell receptor sequencing on B cells isolated from uninjured and post-stroke brains of aged male and female mice. We identified a novel B1b cell progenitor pool distinct from canonical pleural and peritoneal B1 niches. Trajectory analysis showed B1b progenitors transition into age-associated B cell (ABC) subsets, and clonal expansion of IgM+ ABCs (ABC/B1b) and plasma cells following ischemic stroke. We also confirmed analogous ABCs and developing B cell populations in post-mortem human parenchymal tissue isolated from aged brain donors. These studies reveal unique B cell populations that proliferate within the aging CNS and are associated with impaired post-stroke functional recovery in mice. Identification of inflammatory, CNS-resident ABC/B1b cells that are conserved across species is critical as they have the potential to be sequestered from peripheral immunotherapies and/or contribute to age-related neurodegenerative diseases.

## Introduction

Ischemic stroke risk increases with age, and older patients have significantly reduced post-stroke recovery [1, 2]. Rodent models of ischemic stroke replicate the age-related decline in functional recovery attributed, in part, to the aging immune system [3, 4]. Acutely after stroke, there is an influx of innate immune cells in response to damage-associated molecular patterns (DAMPs) originating from CNS cell stress and death [5, 6]. However, during chronic stroke there is a switch to adaptive immune cell infiltration (B and T cells) that persists for months to years after injury [7]. Throughout life, the majority of B cells arise from hematopoietic stem cells in the bone marrow [8], though B1 progenitors also exist in peripheral compartments (e.g. peritoneal and pleural cavities) [9]. B cells progress through developmental stages leading to diverse B cell receptors (BCRs) [9] before leaving the bone marrow and travelling to secondary lymphoid organs, including the spleen and lymph nodes, where they develop into mature B cells and await antigen exposure [8]. Aging induces the additional formation of a Tbet^+^ B cell population [10] known as age-associated B cells (ABCs). ABCs are found in many peripheral compartments and comprise 30-50% of splenic B cells in aged animals. ABCs are memory-like, antigen-experienced, or quiescent B cells that have inflammatory effects depending on disease and tissue context [11]. Despite thorough characterization of ABC phenotypes outside of the CNS, few studies have investigated brain-resident ABCs in aging and after stroke [12].

Previous studies illustrate a detrimental effect of post-stroke B cell populations, including worsened cognitive outcome and CNS-directed autoreactivity [13–15], while others showed a protective role for anti-inflammatory regulatory B cells [16–18]. Remarkably, chronic depletion of CD20^+^ B cell subsets with anti-CD20 antibody after stroke improved cognitive outcomes in young male mice [14] and aged male mice reconstituted with young-male bone marrow also demonstrate improved post-stroke deficits [3]. ABCs specifically induce microglia phagocytosis in male mice, with unclear effects on stroke outcome [12], while previous ABC characterization via single-cell RNA sequencing (scRNAseq) in the periphery only used aged female mice [19]. Of note, oligoclonal bands found in the cerebrospinal fluid (CSF) of stroke patients suggest local antibody synthesis by brain/CSF-resident B cells [20], and patients with elevated CSF antibodies for myelin are at higher risk of post-stroke cognitive decline [21]. Taken together, larger preclinical studies are needed to characterize brain-resident B cell development into ABCs during aging, across sexes, and after ischemic stroke.

To better understand how brain-resident B cells contribute to post-stroke outcomes, we utilized flow cytometry, scRNAseq, and BCR sequencing (scBCRseq) of brain-localized B cells from healthy aged and post-stroke mice. We show that ABC populations expand in both sexes, with novel progenitor B1b cell subsets that directly develop into ABCs with pro-inflammatory phenotypes and expanded clonality after stroke. This highlights a conserved CNS-resident population that may not be exposed to peripheral immunotherapies. Finally, we identify ABC populations in aged post-mortem human brains. Our results reveal that with aging, brain-localized progenitor B cells develop into a proinflammatory ABC phenotype, shared between sexes, that may inhibit recovery after ischemic stroke, with conserved populations present in the aging human brain.

## Methods

All experiments were performed in accordance with the University of Kentucky IACUC under accreditation of AAALAC. Aged (>18 months) male and female C57BL/6JN mice were obtained from the National Institute on Aging. Mice were socially housed and kept in a 14:10 light-dark cycle room with free access to food and water. Methods for tMCAo, behavioral testing, brain processing, B cell isolation, flow cytometry, immunohistochemistry, scRNAseq, scBCRseq, scRNAseq processing & data analysis, high dimensional weighted gene co-expression analysis, transcription factor regulatory network analysis in hdWGCNA, pseduotime trajectory analysis, external data processing, label transfer, and multicompartment aged B cell analysis, clonal analysis, Alpha Fold 3 structural prediction, and human rapid postmortem brain processing are included in the Supplemental Methods document.

## Results

### ABCs are present in aged murine brains and increase after stroke

Flow cytometry on aged post-stroke male and female mouse brains 3 weeks following a 30-min transient middle cerebral artery occlusion (tMCAo) identified B and T cell populations (Supplemental (S)Fig.1), with specific CD11c^+^ and CD11b^+^CD11c^+^ ABCs increasing in all three brain regions (injured cortex, uninjured cortex, cerebellum) after stroke, but by a much larger magnitude in males than in females (Fig.1a,b: statistics in STable 1). A CD23^+^CCR7^+^ B cell memory/activated subset expressing the chemokine receptor CCR7 (facilitating tissue diapedesis) [22], however, exhibited the greatest sex-specific interactions following stroke (Fig.1c). It has been suspected that ABCs may differentiate into autoantibody-secreting cells and transition directly into plasma cells after exposure to CNS antigens [23]. Interestingly, CD138^+^ plasma cells were highest in the aged uninjured female left hemispheres, but also demonstrated significant effects of injury, brain region, and sex interactions (Fig.1d).

**Figure 1:**
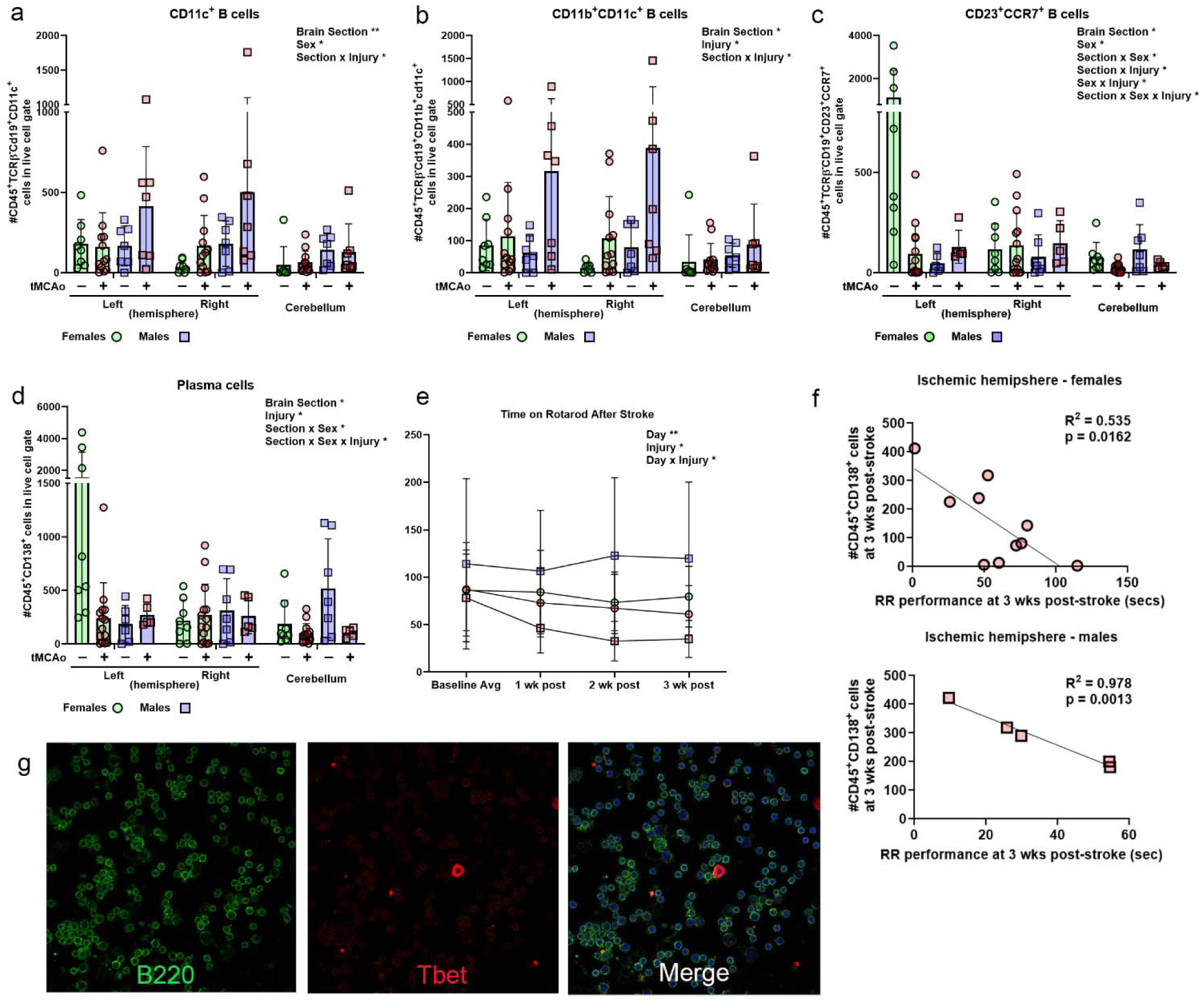
B cell subsets demonstrate sex- and brain area-dependent expression patterns following ischemic stroke that impairs long-term motor recovery. Flow cytometry in immune cells isolated from the left and right hemispheres and cerebellum (designated x axis) in uninjured mice and mice at 3-weeks following a transient middle cerebral artery occlusion (tMCAo; red symbols) show sex-specific interactions for aged females (circles; green bars) vs. males (squares; blue bars). This includes (a) CD11c^+^ and (b) CD11b^+^CD11c^+^ ABCs, with additional interactions for (c) CD23^+^CCR7^+^ memory B cell subsets and (d) plasma cells. (e) tMCAo induced motor deficits in both aged males and females without an effect of sex. Of all B cell subsets, only (f) plasma cells showed a negative correlation with functional recovery in both females (top graph) and males (bottom graph). (g) To confirm expression of the canonical ABC transcription factor Tbet, immunostaining in B cells isolated from the post-stroke brain show high B220 expression (green; left panel) and more reduced Tbet expression (red, middle, right panels) with very large B cell subsets remaining with high Tbet expression.

To identify associations of brain-localized B cell subsets with functional recovery in aging mice, we performed an open field assay to assess anxiety like-behavior, and rotarod assay to assess motor coordination. Open field showed that aged mice, in general, remain in the periphery, indicating anxiety, with no effect of stroke on exploration or anxiety-like behaviors (SFig.1e,f; STable 2). For rotarod, mice were pre-trained for 14 days (SFig.1g) and assessed at baseline and once weekly for 3 weeks post-stroke. Post-stroke aged male mice showed sustained deficits in motor coordination, as reported previously in aged animals [3] (Fig.1e). In both sexes, 3-week motor recovery showed no association with ABC subsets but instead negatively correlated with the number of plasma cells in the ischemic hemisphere (Fig.1f; males R^2^ =0.978, females R^2^ = 0.535). B cells isolated from the post-stroke brain confirmed Tbet^+^ expression [10] in a subset of B220^+^ cells, including larger antibody-producing B cells (Fig.1g). These findings confirm that ABC subsets are present in uninjured aged brains and increase after stroke, with plasma cells, possibly differentiated from ABCs, contributing negatively to functional recovery.

### Single-cell atlas of aged brain-resident B cells reveals developmental stages within the CNS

To gain an in-depth understanding of the heterogeneity of brain-resident B cells in aging and after stroke, we next created a multi-omic single-cell atlas using B cell-enriched single cell suspensions isolated from aged healthy or ischemic brains 3 weeks after tMCAo (Fig.2a). Aged post-stroke males were not included in this study due to the disproportionately high incidence of B cell death that did not meet 10x Genomics requirement for minimum number of live cells per animal. After pre-processing and *in silico* filtering for inclusion of CD45^+^ non-microglia only, 12,081 total cells were obtained from uninjured aged males (n=4, 6073 cells), uninjured aged females (n=4, 3919 cells), and 3-week post-stroke aged female mice (n=4, 2089 cells; SFig.2). CD45^+^ cells contained hematopoietic stem cells (HSCs), monocytes, plasma cells, and a wide diversity of B cells (10,966 cells; CD19^+^/CD20^+^ (*Ms4a1*); Fig.2b,c) spanning multiple developmental stages defined by canonical markers for each population (SFig.2b). This includes *Mki67-*expressing mitotic progenitor B cells and early pro- and pre-B cells (Fig.2d), in line with previously coarsely reported early B cells in the brain [24].

**Figure 2:**
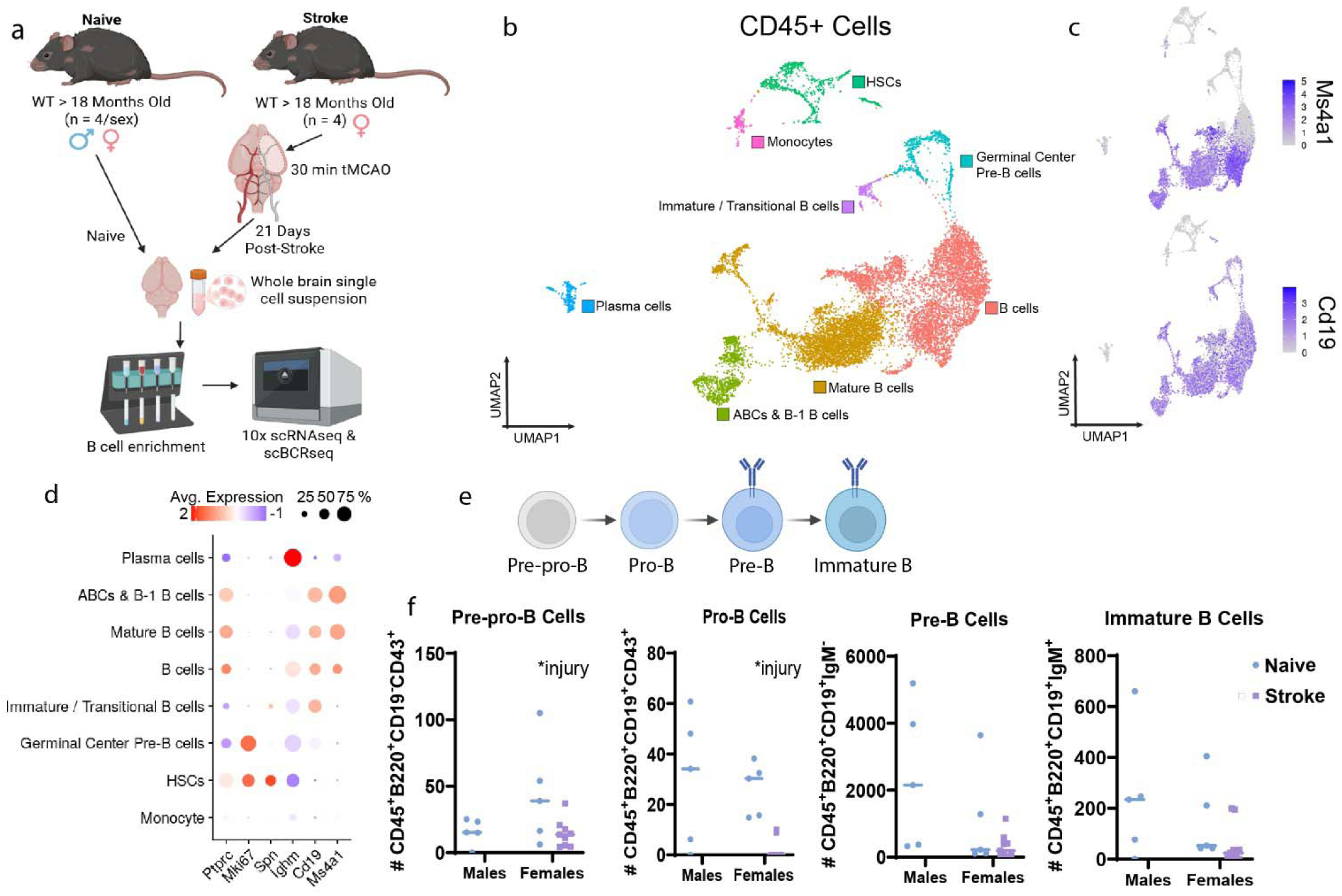
scRNAseq of B cells in healthy aging and after ischemic stroke. **(a)** Experiment design of B cell enriched cell suspension scRNAseq & scBCRseq isolated from healthy aged (Female n = 4; Male n = 4) and 21-day post-stroke mouse brains (Female n = 4; Biorender). **(b)** UMAP of 12,081 total CD45+ cells annotated via canonical marker genes. Hemopoietic stem cells (HSCs; n = 656), monocytes (n = 180), plasma cells (n = 279), all B cells (n = 10,966). **(c)** Gene feature plots of Ms4a1 (Cd20) & Cd19 on all Cd45^+^ cells confirm B cell lineage. **(d)** Dotplot of immature B cell markers across CD45^+^ cells show all developmental stages to age-associated B cell (ABC) populations all unique from the monocyte lineage. **(e)** Schematic of B cell development phenotypes from Biorender. **(f)** Number of immature B cell subsets in uninjured aged mouse cerebellum and at 3wks after transient middle cerebral artery occlusion (tMCAo). Blue circle = naïve for both males (left) and females (right); purple squares = tMCAo.

To expand on the phenotypic resolution of developing B cells (Fig. 2e) and validate our scRNAseq data finding that CNS-resident progenitor B cells are present in aged and post-stroke brains, we identified early developing B cell subsets (Fig.2f; STable 3). To avoid the confounding nature of immune infiltration resulting from blood-brain barrier breakdown in ischemic regions, we performed flow cytometry on the cerebellum, a remote uninjured region of the brain. Pre-pro-, pro-, pre-, and immature B cell populations were present in all groups, with a significant injury effect in pre-pro- and pro-B cell populations (p<0.05; Fig.2g). Thus, the aging CNS houses early developmental B cell subsets creating the potential for local expansion and differentiation within the parenchyma.

### Age-associated B cells develop locally within the CNS

We next reclustered the CNS-resident CD19^+^ population, resulting in 5,835 cells from aged males, 3,646 cells from aged females, and 1,855 cells from post-stroke aged females (SFig.2d-j). B cells were individually labeled via an anchor label transfer with previously published aged mouse B cell annotations (GSE174834) [25] ranging in developmental stage from mitotic B cells to ABCs (Fig.3a; SFig.3). We confirmed these populations with canonical marker genes (Fig.3b) [26]. Cell annotation identified similar population composition shared amongst all three groups, excluding ABCs where post-stroke females had higher mean proportion of ABCs compared to uninjured females (13.4% in post-stroke vs. 4.4% in uninjured; Fig.3c) consistent with our flow cytometry data (Fig.1).

**Figure 3:**
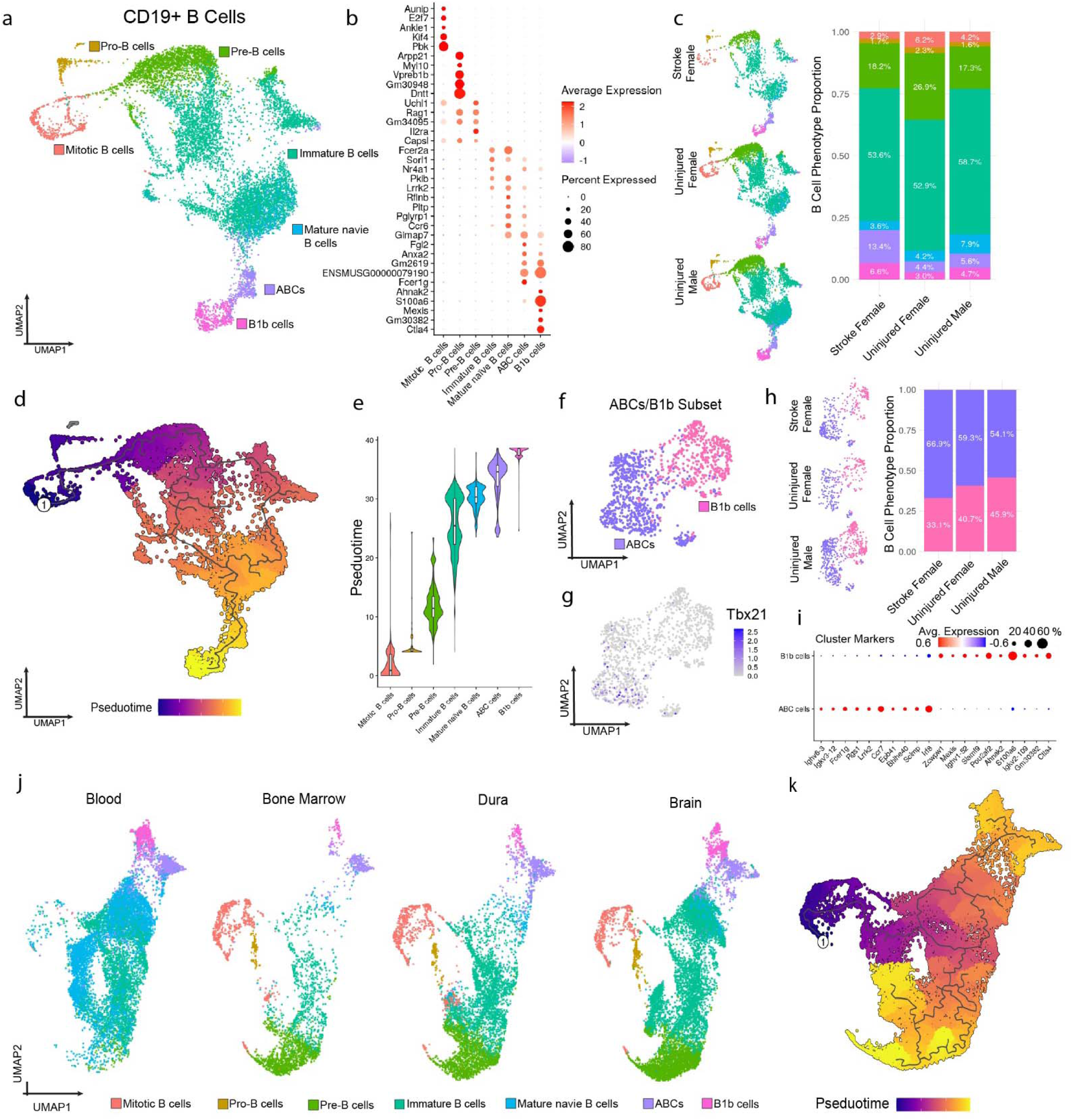
B cells develop into ABCs in the brain and other compartments. **(a)** UMAP of 10,966 brain-resident B cells annotated via canonical marker genes and via an anchor-based label transfer from a published multi-compartment B cell scRNAseq dataset. Mitotic B cells (n = 509), Pro-B cells (n = 203), Pre-B cells (n = 2251), Immature B cells (n = 6141), Mature naïve B cells (n = 659), ABCs (n = 710), B1b cells (n = 493). **(b)** Top 5 B cell population gene markers per identified B cell subset ranked by Log2FC. **(c)** B cell UMAP split by treatment condition (left), with B cell population proportions split by treatment condition and shown in distribution graphs (right). **(d)** UMAP of 10,966 B cells colored Monocole3 pseudotime with trajectories branch tree plotted. Root node labeled by white circled 1 (in dark purple area), determined by identifying earliest branch node inside of *KI67*+ mitotic B cells. **(e)** Violin nested boxplot of pseudotime values across B cell phenotypes ordered increasingly by mean pseudotime**. (f)** UMAP of 1018 ABC/B1b cells colored by phenotype (ABCs, purple; B1b cells, pink). **(g)** Gene feature plot of *Tbx21* (Tbet), the canonical ABC transcription factor. **(h)** ABC/B1b cells split by treatment condition in both UMAP (left) and distribution bar graphs (right). **(i)** Dotplot of top 10 ABCs & B1b cell marker genes (ABCs, bottom row). **(j)** UMAP of a multicompartment integrated 29,806 B cells split across 4 compartments (Blood, Skull Bone Marrow, Dura, Brain) and annotated via identified marker genes to identify B cell subsets. **(k)** Monocole3 pseudotime trajectories of integrated atlas with root node selected from earliest tree node designated as #1 in the dark purple mitotic cell population.

To assess changes across the continuum of B cell development and differentiation, we performed pseudotime trajectory analysis [27] leveraging the known mitotic B cell cluster. This analysis produced a *single trajectory* from mitotic B cells, traversing pro- and pre-B cells, through immature and mature stages to finally culminate in ABCs (Fig.3d,e). These results demonstrate a continuum of B cell development in the aging brain culminating in transcriptionally unique ABC/B1b cells that are significantly enriched in the 3-week post-stroke period.

To better understand this novel ABC/B1b phenotype, we reclustered these cells and obtained two resulting populations (Fig.3f). The ABC cluster, enriched for *Tbx21* (Fig.3g), comprises a greater proportion in post-stroke females compared to uninjured females (Fig.3h). Cluster markers outline genes that differentiate ABCs from B1b cells (Fig.3i; SFig.4). To confirm terminal differentiation into an ABC phenotype, we integrated a publicly available dataset with B cells from aged mouse blood, skull bone marrow, and dura [25] with our own brain data (Fig.3j; SFig 5a-c). CNS-resident B cell composition mirrors the composition of skull bone marrow and dura B cells, namely via a large population of pre-B cells (SFig 5d-f) compared to blood. Pseudotime trajectories of combined compartments show the same population of ABCs at the end of one trajectory as found in the aged brain (Fig.3k). Furthermore, expression of *Fgl2* (fibrinogen-like protein 2), a consistent cluster marker of ABCs, represents a novel marker of mouse ABCs irrespective of compartment (SFig.5g).

### ABCs exhibit activated phenotypes

Differential gene expression (DEG) identified gene changes secondary to stroke (uninjured vs. post-stroke females) or with aging in uninjured male vs. female mice (Supplemental Data Set 1). Post-stroke females compared to uninjured females showed 519 genes (up: 471; down: 48) to be differentially expressed after stroke (Fig.4a) while aged mice demonstrated 593 genes (up: 430; down: 163) differentially expressed between uninjured males versus females (Fig.4b). DEGs after stroke within female mice and sex differences with aging resulted in an overlap of 228 (33.9%) upregulated genes and 40 (23.4%) downregulated genes, with 83% downregulated genes after stroke shared with aging males (Fig.4c). Biological process gene ontology (GO) analysis of upregulated genes after stroke revealed terms enriched for cytoplasmic translation, mature B cell differentiation, cytokine production involved in inflammatory response, and positive regulation of B cell proliferation (SFig.6a). Biological process gene ontology of stroke upregulated genes was very similar to GO terms seen in between males and females in healthy aging, with terms enriched for increased cytoplasmic translation, protein metabolism, positive regulation of cytokine production, both suggesting increased metabolic activity and activation in response to inflammation. Taken together, these results suggest a link between brain-resident B cell activation responses of stroke-induced inflammation and sex differences in aging, with high pro-inflammatory markers in males even in the uninjured state.

**Figure 4:**
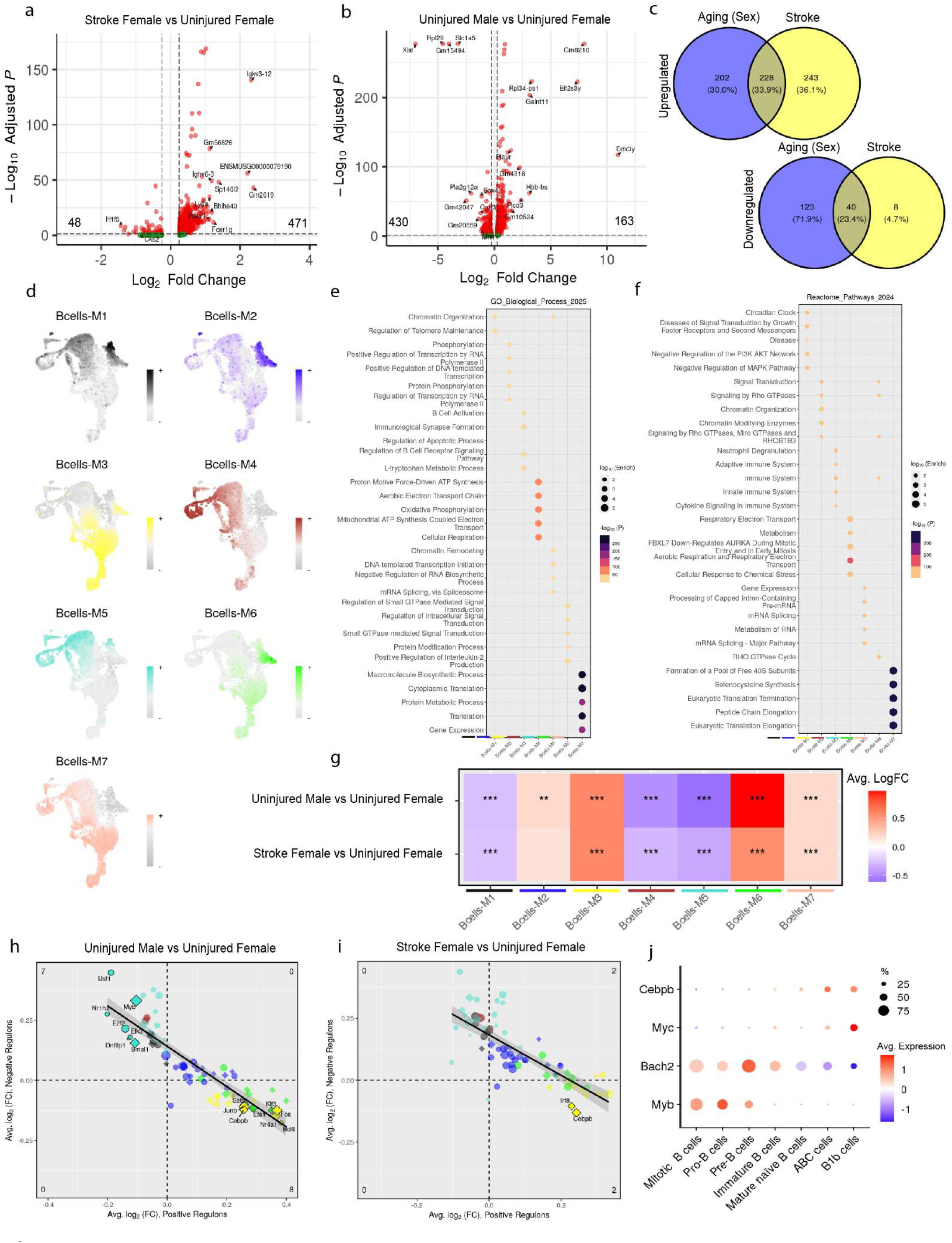
Gene and transcription factor regulator networks reveal shared inflammatory signaling between uninjured male and post-stroke female B cells (a-b) Volcano plots of differentially expressed genes between treatment conditions (Log2FC > 0.25 or < -0.25; Padj < 0.05, min.pct > 0.1; Top 10 genes by Log2FC labeled). **(c)** Venn diagram of significant up- and downregulated genes shared between stroke female vs uninjured female and uninjured male vs uninjured female. **(d)** hdWGCNA Module expression score feature plots of first 1000 genes in each module on 10,966 brain resident B cells. **(e)** Top 5 significant Gene Ontology (GO) Biological Process 2025 terms for each module via Enrichr. Dot size is Enrichr enrichment score and colored by -Log10 adjusted P value. **(f)** Top 5 significant Reactome Pathway 2024 terms for each module via Enrichr. Dot size is Enrichr enrichment score and colored by -Log10 adjusted P value. **(g)** hdWGCNA differential module score heatmap (Log2FC > 0.2 or < -0.2, Padj < 0.05, Wilcox rank sum, pseudocount = 0.01; ** Padj < 0.01, *** Padj, < 0.001). **(h-i)** Transcription Factor (TF) differential regulon scatter plot across treatment conditions where predicted positive relationship regulons (promoting) plotted on the X-axis and predicted negative relationship (suppressing) regulons plotted on the Y-axis. TFs are colored by module association (See hdWGCNA methods for more detail). **(j)** Dotplot of *Myb*, *Bach2*, *Myc*, *Cebpb* across B cell phenotypes.

We utilized high dimensional Weighted Gene Co-expression Analysis (hdWGCNA) [28] to create correlated gene expression modules to investigate gene-specific changes across the B cell developmental trajectory. After dynamic tree splitting, 7 modules were present that represented both B cell population-localized and diffuse gene expression networks (Fig.4d; SFig. 6b,c). Interrogation of module-specific functionality revealed modules significantly enriched for B cell activation and proliferation (M3), chromosome reorganization (M1, M5), cellular respiration and mitochondrial activity (M4), Rho GTPase activity (M2, M6), and ribosomal genes (M7; Fig.4e,f). Genes encompassed within M3 relating to B cell activity were highly enriched in mature B cells (*Stap-1, Fcrl1, Gpr183*) and ABCs (*Cd81, Cd22, Themis2*; SFig.6d). We examined CNS-resident B cell module enrichment across stroke and aging conditions by computing differential module expression scores, utilizing the top 1000 genes in each module. Module M3, containing genes associated with B cell activation and BCR signaling and enriched in mature B cells and ABCs, was significantly upregulated in post-stroke vs. uninjured females (Log_2_FC=0.583; P_adj_=5.19E-45) as well as in uninjured males vs. uninjured females (Log_2_FC=0.588; P_adj_=2.68E-81). The same pattern of upregulation can be seen in M6 with stroke (Log_2_FC=0.564; P_adj_=2.79E-16) and in uninjured males (Log_2_FC=0.969; P_adj_=2.67E-106; Fig.4g). M6 is also enriched in mature B cells and ABCs. M7, albeit not B cell lineage-specific, demonstrates a conserved global increase in GTPase motility and gene expression similarly upregulated with stroke and in uninjured males. Utilizing Rank-Rank Hygrometric Overlap, we found a significantly strong correlation (r=0.64; p=2.2*10^-16^) between transcriptomic profiles of post-stroke females vs. uninjured females and uninjured male vs. uninjured female (Fig.4g; SFig.6e,f). These results suggest that this transcriptional signature, shared between stroke and aging sex differences and seen in coarse DEG analysis, is related to B cell activation in mature B cell and ABC/B1b CNS-localized populations.

### Transcription factor regulation network analysis implicates *Cebpb* regulon driving ABC activation

To understand if the B cell activation transcriptional signature is driven by specific transcription factors (TF), we further utilized hdWGCNA [28] to investigate the transcriptional programs within the identified modules by utilizing machine learning models to infer regulatory relationships between TFs and their predicted targets [29, 30]. The increase in B cell activation in uninjured aged males over uninjured aged females (module M3) is driven by a TF regulon ensemble encompassing *Cebpb, Junb, Fos, Nr4a1*, and *Bcl6.* These TFs are not only differentially expressed, excluding *Bcl6*, but their positive target genes are upregulated and differentially expressed in uninjured male B cells compared to females (Fig.4h; SFig.7). Comparing differential TF regulons between post-stroke females and uninjured females reveals *Cebpb* and *Irf8* (Fig.4i; SFig.7f). Their target genes are also differentially expressed in post-stroke females relative to uninjured females, specifically – and again – in B cell activation module M3. *Cebpb* was previously implicated in a B cell phagocytic phenotype after injury [31]. These findings outline a potential TF program that drives sex-specific B cell activation via a shared TF, *Cebpb*, that is also activated after stroke in females.

We calculated hdWGCNA module pseudotime trajectories to investigate how the modules changed within our developmental trajectory. Modules M3, M6, and M7, previously identified to be rich in B cell activation genes, have the highest module expression scores in the ABC/B1b population (SFig.8a), suggesting increased B cell activation in ABCs after stroke and in uninjured males compared to uninjured females. We also investigated if any previously identified TFs were significantly localized across pseudotime, with a specific focus on mature B cells and ABC/B1b cells. Investigation of TF network dynamics across differentiation demonstrates that early B cell populations are *Myb*^+^ & *Bach2*^high^ and, during transition, are marked by a loss of *Myb*, previously shown to induce Tbet (*Tbx21*) after its loss while decreasing *Bach2* [32]. Concurrently, the induction of *Myc* and *Cebpb* occurs in mature and late-stage ABC/B1b populations (Fig.4j), suggesting the development into a phagocytic phenotype [24]. We further confirmed the previous observation of B cell development phenotype-specific TF recruitment across compartments *of Myb, Bach2, Myc* and *Cebpb* (SFig.8c). These findings implicate multiple TFs in the concerted activation of mature B cells and ABCs in the brain.

### Stroke induces clonal expansion of IgM^+^ B cells in the aged brain

We performed concurrent scBCRseq to investigate the effects of aging and stroke on brain-resident BCR repertoires. Density plots laid over the initial UMAP with all CD45^+^ cell populations confirm BCRs are present in our identified B cell clusters across all groups (Fig.5a). We defined BCR clones by identical VDJ gene usage in both heavy and light chains, allowing for amino acid changes and somatic hypermutation. Hyperexpanded clones (defined as comprising greater than 10% of the total BCR repertoire) were found primarily in the ABC/B1b and plasma cell clusters (Fig.5b). Clonal network analysis identified BCR sequence similarity amongst cell populations using either plasma cells or ABC/B1b cells (Fig.5c) as the starting node. Both the plasma cell cluster and ABC/B1b cell cluster share clonal similarity with other B cell developmental populations, again indicating the potential for CNS-specific proliferation. Post-stroke females exhibited the largest proportion of hyperexpanded clones, while uninjured males had no hyperexpansion (Fig.5d,e). Shannon index scores confirmed BCR clonal diversity [33], in which a low score indicates decreased diversity presumably due to clonal expansion. Uninjured females had a Shannon index of 4.12 vs. post-stroke females (4.00; Fig.5f), with the differences between groups largely driven by one mouse. Interestingly, in all groups, the predominant heavy chain constant region was IgM (Fig.5g), indicating minimal class switching. Amongst the hyperexpanded clonotypes, *IGHV7-3* was the most expressed variable gene in both post-stroke and uninjured females (Fig.5h).

**Figure 5:**
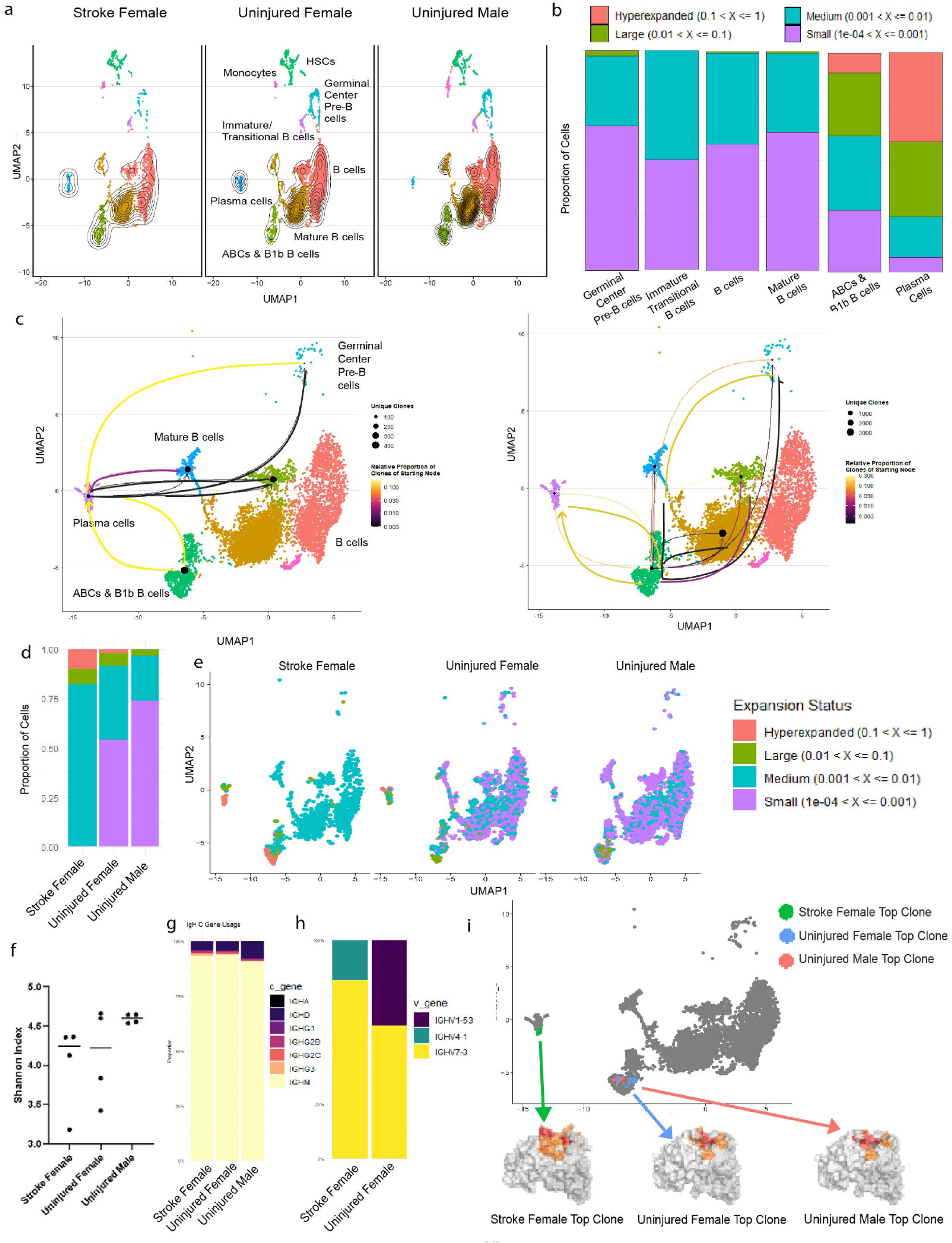
Stroke induces hyperexpansion of IgM^+^ ABC/B1b B cells and plasma cells. **(a)** UMAP of B cell clusters overlayed with BCR density contour, confirming sequenced BCRs were present in the expected populations across groups. **(b)** Clone size across the identified B cell subtypes. **(c)** BCR clonal network analysis showing the network with plasma cells as the primary node (left) and the ABC/B1 cluster as the primary node (right). **(d)** Clonal expansion status across groups. Injured females showed the highest proportion of hyperexpanded cells, followed by uninjured females. **(e)** UMAP of B cells showing expansion status across. **(f)** Shannon diversity indices show minimal difference in diversity across groups. **(g)** IgH constant chain usage is large IgM across groups. **(h)** variable gene usage of hyperexpanded clones in uninjured and post-stroke females. Uninjured males did not have any hyperexpanded clones. **(i)** top expanded clone in each group depicted on UMAP of B cells. post-stroke female top clone is in the plasma cell cluster while uninjured males and females are ABC/B1b cluster. Structural predictions of binding regions are depicted. Red region is 0.45pt confidence, orange region is 0.40pt confidence.

We generated ABodyBuilder2 Fv models for the six representative antibodies (top two clonotypes per group) and predicted paratope residues with proABC-2 (Fig.5i). The top clone in uninjured animals of both sexes was in the ABC/B1b cell cluster, while the top post-stroke clone was in the plasma cell cluster (Fig.5i). Using a conservative cutoff (pt ≥ 0.45), antibodies contained a median of 6.5 predicted contact residues (range 3–7), with most contacts on the heavy chain (median 4 residues; range 3–5). At the sensitivity cutoff (pt ≥ 0.40), the predicted interface expanded to a median of 18.5 residues (range 18–22) while remaining strongly complimentary-determining region (CDR)-enriched (median 89.2% of residues in CDRs). Predicted contacts consistently involved CDR-H3 (positions 95–102; present in all six antibodies), which is the largest portion of the binding surface and is key to determining antibody specificity, whereas no CDR-H1 residues exceeded pt ≥ 0.45. Residues above each threshold were mapped onto the Fv structures to visualize candidate binding surfaces for future antigen-focused experiments.

### Identification of ABC populations in human post-mortem brains

The presence of brain-localized ABCs was confirmed in humans who were participants in the University of Kentucky Alzheimer’s Disease Research Center (UK-ADRC) autopsy cohort. Details of the recruitment, inclusion/exclusion criteria, and follow-up from the UK-ADRC cohort have been published previously [34, 35]. An emphasis of this cohort is on the rapid post-mortem (RPM) biobank, with tissue collection near time of death (autopsies usually performed ∼2-3 hours postmortem). Samples of left and right motor cortex and cerebellum (lacking dura) were collected from 14 RPM donors with an average age of death at 86 years (Fig.6a; STable 4). Among the included participants, 77% (n=10) had cognitive impairment documented before death, with 69% having the diagnosis of dementia (n=9). Three study donors (23%) had a documented prior history of stroke. Results of left and right brain samples were averaged after confirming no cell population differences between hemispheres. Both CD11b^+^ and CD11c^+^ ABCs (Fig. 6b-d) were broadly present within the CD19^+^ B cell populations isolated from every brain region, which also showed high CD23^+^ expression (Fig.6e). The proportion of CD11b^+^ ABCs in both the cerebellum and motor cortex showed no significant sex differences, though males tended to have a lower proportion than females in the cerebellum (p=0.093; STable 5). CD11c^+^ ABCs comprised a wide range, with proportions ranging from 2-75% of B cells. There were no significant differences detected in ABC proportion between sexes in either location, though males exhibited a significant increase in cerebellar CD11c^+^ ABCs with increasing age, a correlation not found within the same RPM donors for the CD11b^+^ ABCs (Fig.6f,g). The 3 donors with prior stroke exhibited some of the lowest representative values for CD11c^+^ ABCs. We confirmed ABC presence in one of the RPM brain donors with prior stroke (82-year-old male) using immunohistochemistry to confirm CD11c^+^ ABCs in parenchyma and not in unperfused vasculature or leptomeninges (Fig.6h). Investigation of a scRNAseq dataset of human brain and leptomeningeal B cells (GSE288856) [36] in Alzheimer’s Disease (AD) verified that CD11c^+^ (*Itgax*) T-Bet^+^ (*Tbx21*) ABCs are proportionally enriched in AD patients (40%) compared to case controls (18%), along with phagocytic and immature B cell populations (Fig.6i).

**Figure 6:**
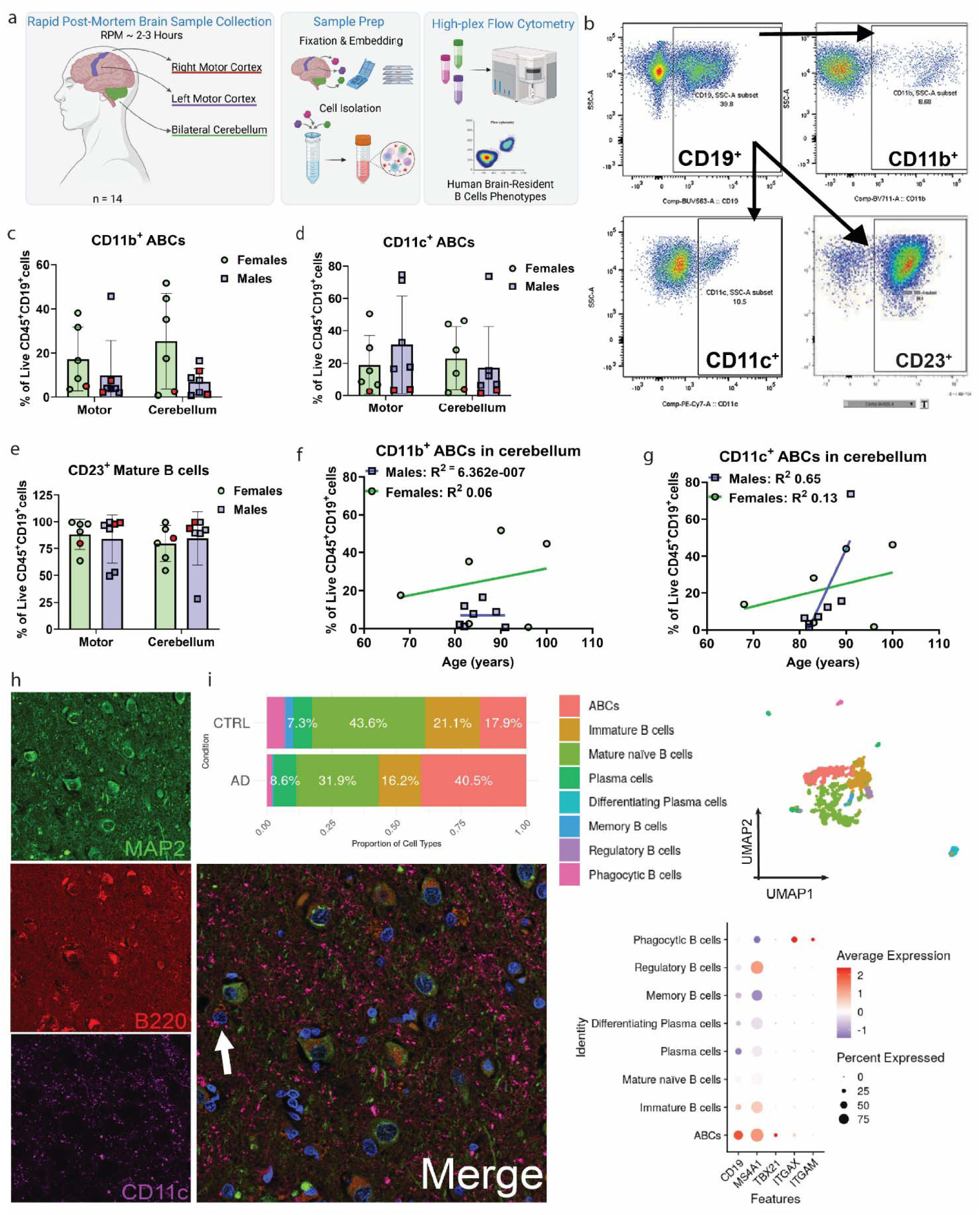
ABCs are present in human rapid post-mortem post-stroke brain. **(a)** Experimental overview for collection of rapid post-mortem human brain samples. **(b)** Gating strategy for flow cytometry of human brain samples. Samples were first gated for time, single cells, and live cells (not shown). **(c-e)** CD11b^+^ **(c)** and CD11c^+^ **(d)** ABCs and CD23^+^ mature B cells **(e)** in human motor cortex and cerebellum expressed as percent of live CD45^+^CD19^+^ cells. Females = green circles. Males = blue squares. Red depicts donors with a prior reported history of stroke. **(f-g)** correlation of age with percentage of CD11b and CD11c ABCs in cerebellar tissue show male-only increase with age for CD11c^+^ ABCs. **(h)** Immunohistochemistry staining of post-mortem human brain for MAP2 (green, neurons), B220 (red), CD11c (purple) identifies B220^+^ CD11c^+^ MAP2^-^ B cells (arrow). **(i)** UMAP of human B cells from GSE288856 of brain and leptomeninges with proportion of cell types grouped by case diagnosis of Alzheimer’s disease (AD) vs. cognitively normal control (CTRL). Dotplot of B cell and ABC markers across B cell subsets for the B cell-related genes *Cd19*, *MS4A1*, *Tbx21, ITGAX*, and *ITGAM*.

## Discussion

We identified that stroke, in the context of aging, results in an expansion of pro-inflammatory ABCs, and that the aged brain houses early parenchymal-based B cell progenitors. Trajectory analysis demonstrated a continuum of B cell development from these early progenitors through immature and mature subsets to terminally differentiate into the stroke-expanded *Bach2*^low^ *Cebpb*^+^ ABC/B1b phenotype. We corroborated the finding of ABCs in human cortical and cerebellar tissue, identifying populations that are specifically affected by sex and age. If there is an early developmentally-derived CNS B cell pool maintained by local progenitors as our study supports, treatments can be developed to specifically target this niche, particularly in aged individuals with impaired peripheral B cell development.

The *de novo* development of B1 cells is considered to occur from the yolk sac (first wave) and fetal liver (second wave and predominant source) prior to bone marrow (third wave), though populations are largely replenished with age through self-renewal driven by Bach2 [37]. Of critical note, IgM^+^ B1 cells canonically reside in pleural and peritoneal cavities, and the presence of Bach2 is a survival factor for B1 cells in peripheral niches [9, 38]. To our knowledge, B1 cells have not been studied in the brain. Interestingly, B1a and B1b B cells are increased in the cervical lymph nodes of 3xTgAD Alzheimer phenotype disease mice [39], and innate-like marginal zone B cells play a role in post-stroke infection prevention [40]. More recent work identified early proliferating B cells in young brain, leptomeninges, and dura, with brain expansion in the acute disease phase of experimental autoimmune encephalomyelitis (EAE), a preclinical model of multiple sclerosis [41]. However, our data are the first to localize the development of these functionally-relevant innate-like B cells to the CNS in the aged brain (>18 mos. old) and after ischemic stroke.

ABCs are memory-like or antigen-experienced B cells which have been most extensively studied in models of autoimmune diseases like multiple sclerosis and lupus [11]. Little is known about their role(s) in the healthy aged brain and after stroke [12], though a previous study showed that CD11b^high^ ABCs induce changes in microglia that result in higher rates of microglial phagocytosis 7 days post-stroke [12]. We, however, are the first to show that these B cells may terminally differentiate locally in the brain as opposed to the periphery. This is significant as it implicates the brain as a functioning ectopic lymphoid tissue both in normal aging and after stroke, supporting sustained B cell development. Further studies should determine if enhanced post-stroke differentiation requires the help of T cells, as prior studies have shown [14], or if this is a T cell-independent process more akin to B1 maturation. Interestingly, our flow cytometry data show that these populations increase in both hemispheres after stroke, not only the ischemic hemisphere, a finding in line with our prior studies that identified increased B cell populations remote from the infarct that support neurorecovery [16, 42].

Meningeal B cells originate predominantly from the calvaria bone marrow and can enter the brain directly, bypassing systemic circulation [43]. We are limited to conclude that early immature B cells divide and transition through maturity *exclusively* within the brain and do not migrate from the periphery or dura. However, BCR analysis shows expansion of IgM^+^ brain-resident mature B cells, ABCs, and plasma cells, suggesting a within-brain transition for all B cell subsets. Additionally, we found a strong negative correlation between total plasma cells in the CNS and post-stroke motor performance. Antibody-producing B cells contribute to post-stroke cognitive impairment in preclinical models [14], with IgM antibodies remaining in the brain of stroke patients for years [14]. Interestingly, our work did not identify class-switched hyperexpanded IgG or IgA BCRs, which confirms similarity to the polyreactive, nonspecific IgM^+^ autoantibodies traditionally formed by B1b cells. Clinical data show a detrimental association of self-reactive antibodies and cognitive decline [44]. It remains unknown what the antigen targets are for the expanded clones identified in this study; future studies will be required to elucidate possible B cell-derived CNS autoantibodies and how they may hinder stroke recovery. But of note, only B cells from females were hyperexpanded and may contribute to brain-localized plasma antibody-producing populations though the caveat remains we did not scBCRseq post-stroke males.

Male post-stroke mice exhibit the greatest increases in ABC populations in flow cytometry data. However, as mentioned we could not isolate sufficient live male post-stroke B cells at 3 weeks post-stroke to perform scRNAseq in this group, highlighting a limitation of this study. Advances in single cell sequencing technologies that allow for sequencing of fixed whole cells may overcome this technical hurdle [45]. Large sequencing efforts in other brain-resident immune cells, such as microglia, demonstrate similar age-associated sex differences in activation of male immune cells [46]. This transcriptional signature of activation was also upregulated after stroke in females in ABCs and, after TF analysis, was inferred to be controlled by *Cebpb. Cebpb* was previously identified to be increased in peripheral blood mononuclear cells of elderly patients [47] and has been implicated in inducing post-injury B cells into a myeloid like phagocytic phenotype [48], consistent with previously described ABC function [49]. Future studies that directly perturb B cell *Cebpb* in aged rodent models are required to elucidate how *Cebpb* and its target genes work to increase B cell activation, specifically in ABCs, and their functional relevance to health and recovery in the aging brain.

Our postmortem data confirmed the presence of both CD11b^+^ and CD11c^+^ ABC populations in the human brain, with a range of expression up to 80% of total CD19^+^ B cells. Most B cells also expressed CD23, which is associated with mature and activated B cells [50]. We validated these populations in a publicly available human scRNAseq data set to show a significant ABC enrichment in Alzheimer’s disease. We found gender-specific and brain location-specific differences in ABC populations, with caveats related to sample sizes that limited statistical power. We also noted that CD11b^+^ ABCs trended to be higher in females when isolated from the cerebellum. CD11b^+^ ABCs in healthy human donor blood exhibit migratory properties [51], and CD11b facilitates B cell migration by aiding in adhesion to endo- and sub-endothelium in the vasculature via the intercellular adhesion molecule 1(ICAM1) [52]. These peripheral B cells express CD11b, regardless of chemokine presence and attraction, and also expressed CD27, a memory B cell marker [52]. Future studies should include CD27, CCR7, and CXCR5, the latter receptors instrumental in homing for CNS-specific migration from parenchymal ectopic lymphoid structures [53], to confirm migratory versus CNS-localized progenitor B cell pools in aged humans.

In conclusion, this study is the first to identify a B cell resident lineage in the aged brain that develops from early progenitors to ABCs. We also show that aged male mice have a substantially increased baseline B cell inflammatory state in the brain over female counterparts. Stroke induced clonal expansion of IgM^+^ ABC/B1b and plasma cells that exhibit a phagocytic gene signature. We also confirmed developing, mature and ABC B cell subsets in human parenchyma. These findings have important implications for future work examining sex differences, brain injuries, and neurodegenerative diseases in the aging brain. Stroke, traumatic brain injury, and Alzheimer’s disease have all had clinical trials with peripheral immunotherapies that have shown a broad range of results in terms of efficacy and safety. It is possible that detrimental – or perhaps even some beneficial – immune effects originate from CNS-localized, self-renewing B cell pools sequestered from peripheral immunotherapies and thus may require unique CNS targeting for therapeutic efficacy in diseases of aging.

## Acknowledgments

Thank you to the brain donors and their supportive families for their invaluable contribution to this work. These studies are supported by grants from the National Institutes of Health to AMS: R01NS088555, RF1NS088555; AMM: F30NS141486; MKC: T32AG057461; TU: T32NS077889, 3RF1NS088555-07A1S1; KMC: T32NS077889; SJC: R01NS129668; UK-ADRC: P30 AG072946. Services and support for scRNAseq experiments were provided by the College of Arts & Sciences Imaging Center at the University of Kentucky. This work was supported by the UK Light Microscopy core facility (RRID: SCR_026405), Flow Cytometry & Immune Monitoring core (RRID: SCR_026358) and Biospecimen Procurement & Translational Pathology Shared Resource Facility (P30CA177558).

## Data Availability

Publicly available B cell sequencing data was obtained from GSE174834. Raw and Processed data from this study will be available upon publication and deposit on a publicly available site with link to ODC-Stroke.

