## Supplemental Tables and Figures for "Age-associated B cells (ABCs) develop from a CNS-localized progenitor pool into a pro-inflammatory phenotype after stroke"

*equal contribution

**Supplemental Data Figures and Tables
Supplemental Figures**

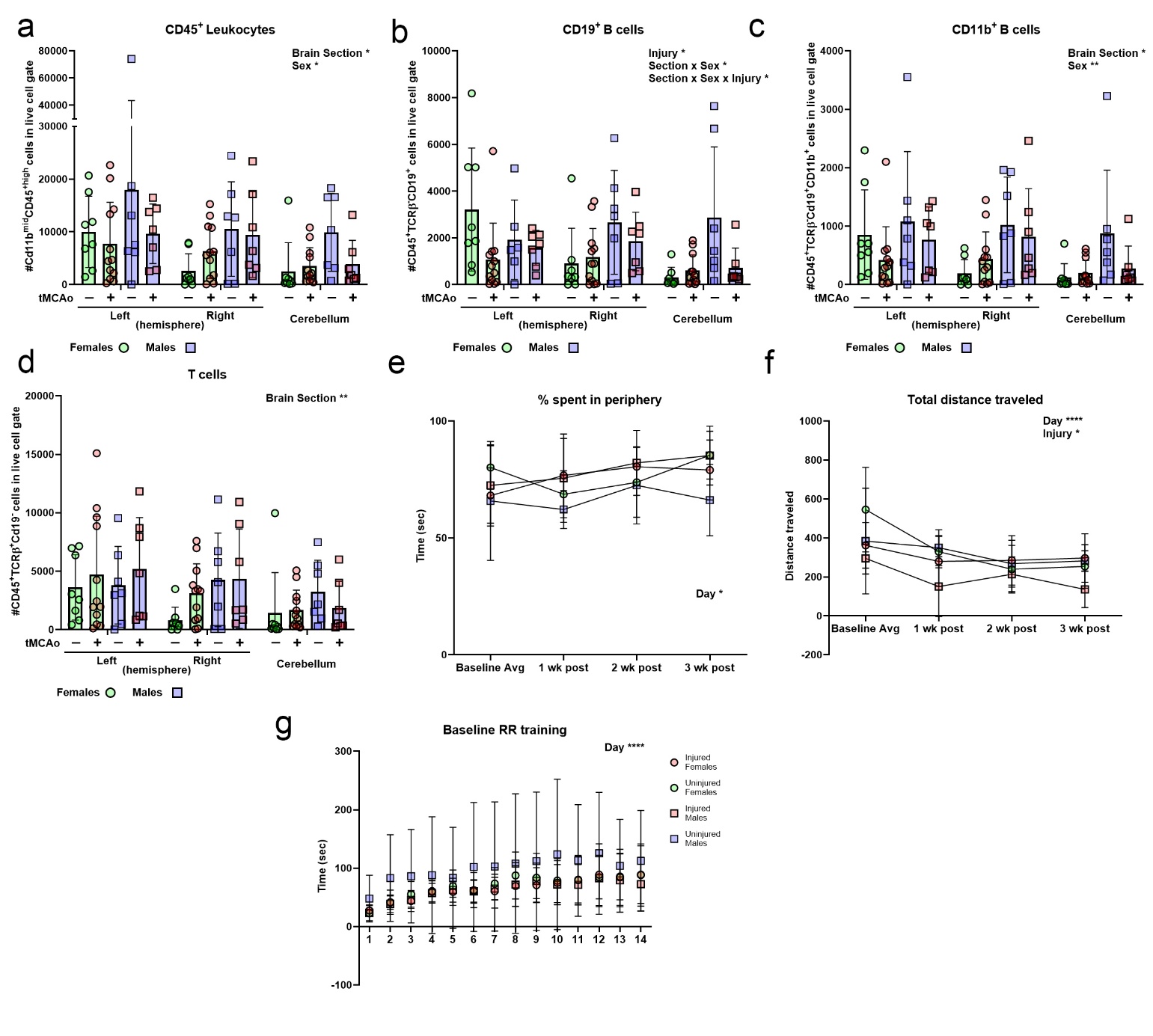
**SFigure 1: B cell subsets demonstrate sex- and brain area-dependent expression patterns following stroke**

Flow cytometry in immune cells isolated from the left and right hemispheres and cerebellum (designated x axis) in uninjured mice and mice at 3-weeks following a transient middle cerebral artery occlusion (tMCAo; red symbols) show sex-specific interactions for aged females (circles; green bars) vs. males (squares; blue bars). This includes **(a)** CD45^+^ general leukocytes and **(b)** CD19^+^ B cells, with additional interactions for the **(c)** CD11b^+^ ABC subset but not **(d)** T cells. **(e)** tMCAo had mild effect on peripheral localization in the open field for both aged males and females without an effect of sex, though the stroke **(f)** did reduce the distance traveled in the open field for all mice. **(g)** All mice trained equally without sex-based differences for 10 days on the rotarod prior to randomization to cohort.

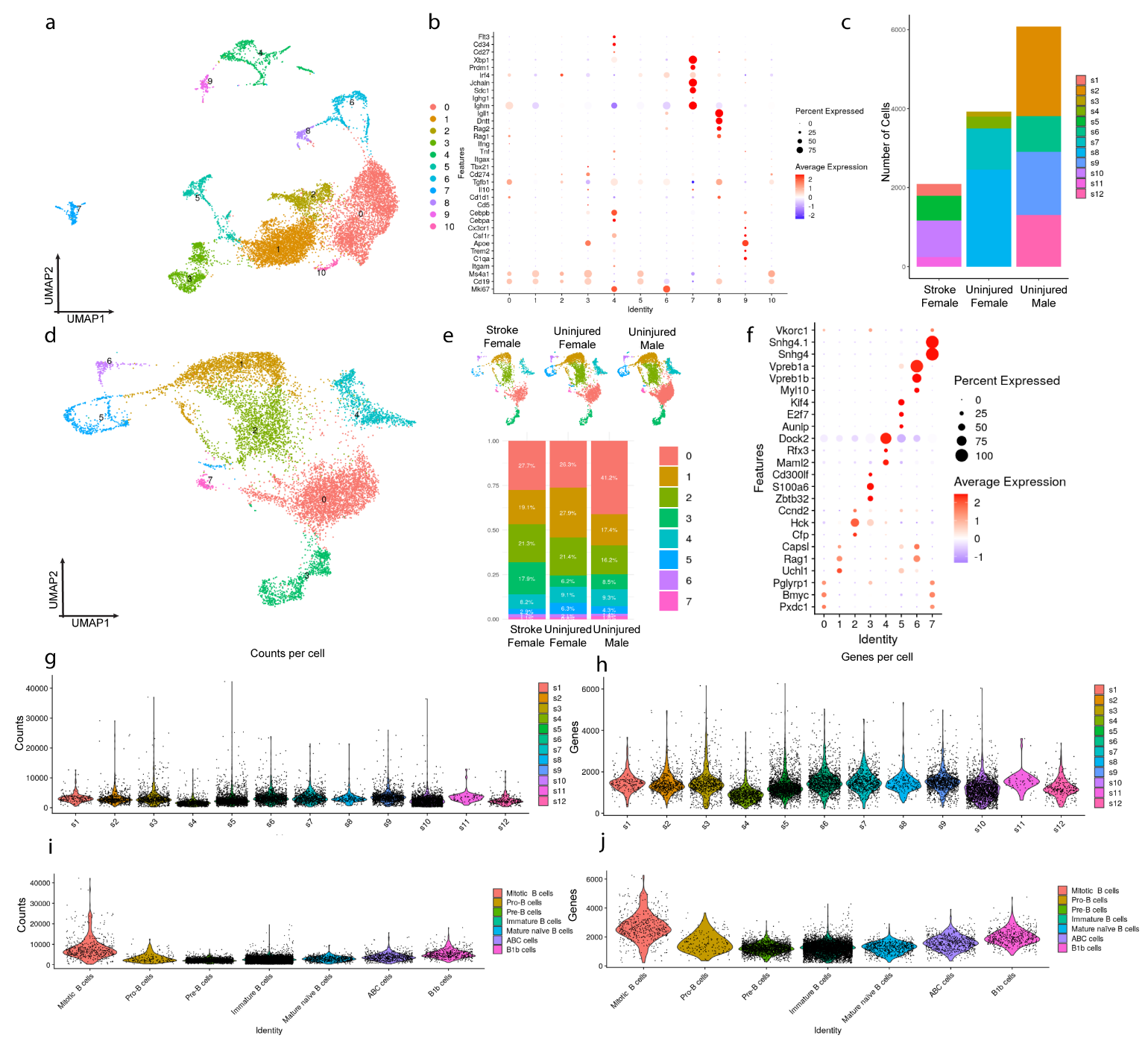

**SFigure 2: scRNAseq quality metrics**

**(a)** UMAP of 12,081 total CD45^+^ cells annotated via clusters at resolution of 0.2. **(b)** B cell phenotype gene marker dot plot shows initial B cell subsets. **(c)** Stacked Boxplot of number of cells per sample per treatment group. **(d)** UMAP of 10,966 brain-resident B cells annotated via clusters at resolution of 0.2 **(e)** B cell UMAP split by treatment condition (top). B cell population proportions split by treatment condition (bottom) for the 7 finalized subsets. **(f)** Top 3 B cell cluster gene markers ranked by Log2FC for the 7 subsets. Violin plot of **(g)** counts per cell grouped by sample; **(h)** features/genes per cell grouped sample; **(i)** counts per cell grouped by B cell phenotype; and **(j)** features/genes per cell grouped by B cell phenotype.

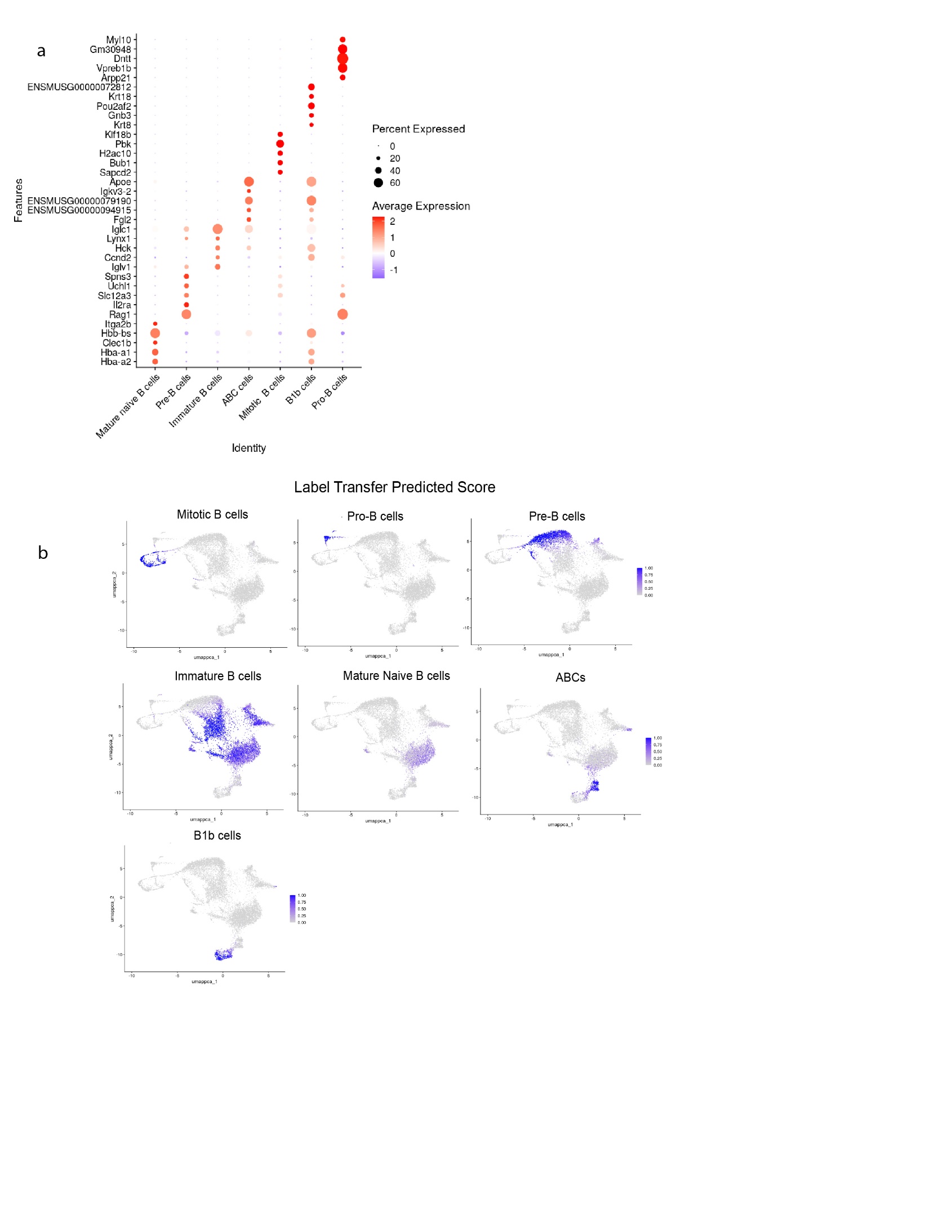

**SFigure 3: Production of B cell annotation dataset**

**(a)** Top 5 cluster markers of B cell phenotypes ranked by Log2FC. **(b)** Label transfer prediction score of B cell phenotypes anchor based label transfer onto brain derived B cells**.**

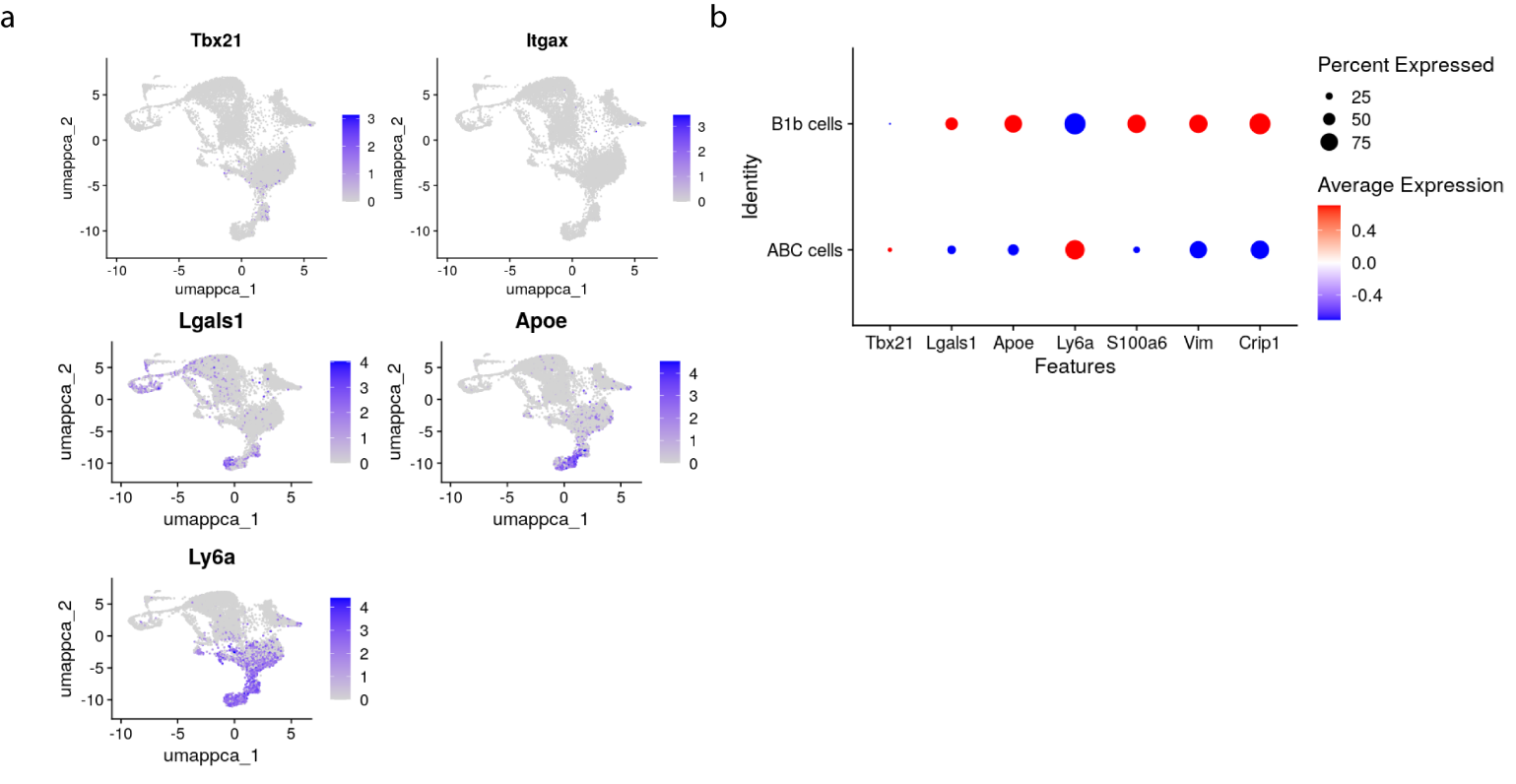

**SFigure 4: Characterization of ABC/B1b cells**

**(a)** Gene feature plots of ABC/B1b markers *Tbx21* (Tbet), *Itgax*, *Lgals2*, *Apoe* ,and *Ly6a*. **(b)** Dot plots of ABC/B1b markers *Tbx21* (Tbet), *Itgax*, *Lgals2*, *Apoe* ,*Ly6a* characterize the overall expression patterns between the two subsets at the terminal point of the pseudotime trajectory.

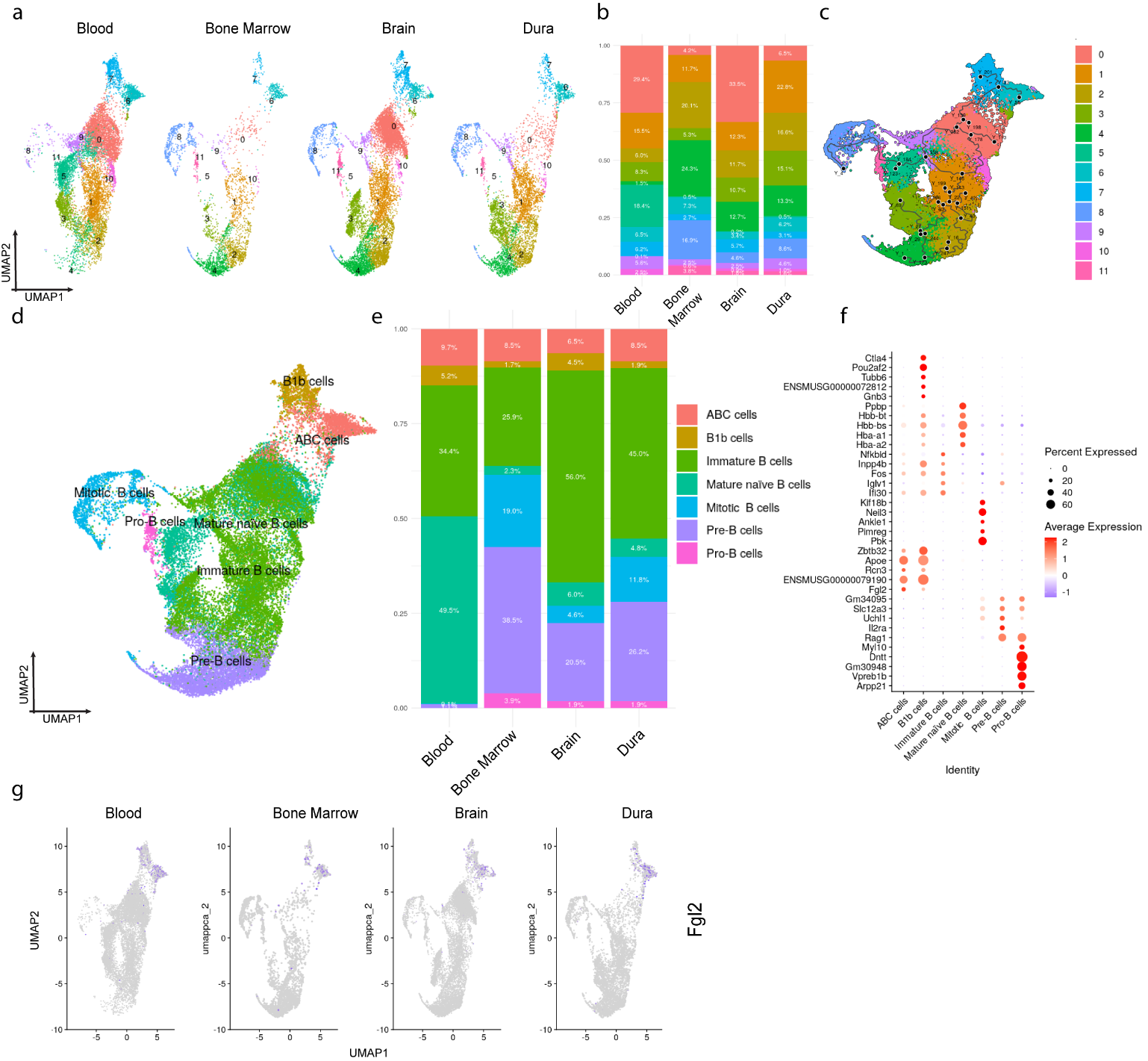

**SFigure 5: Integrated multicompartment B cell dataset with brain-resident B cells**

**(a)** UMAP of 29806 total B cells integrated across 4 compartments annotated via clusters at resolution of 0.2. **(b)** Cluster proportions across compartments. **(c)** Monocole3 pseudotime across clusters. **(d)** B cell phenotypes of integrated multi compartment atlas. **(e)** B cell phenotypes proportions across compartments. **(f)** Top 5 B cell phenotype cluster markers ranked by Log2FC. **(g)** Gene feature plot of *Fgl2* plotted on B cell atlas split by compartment.

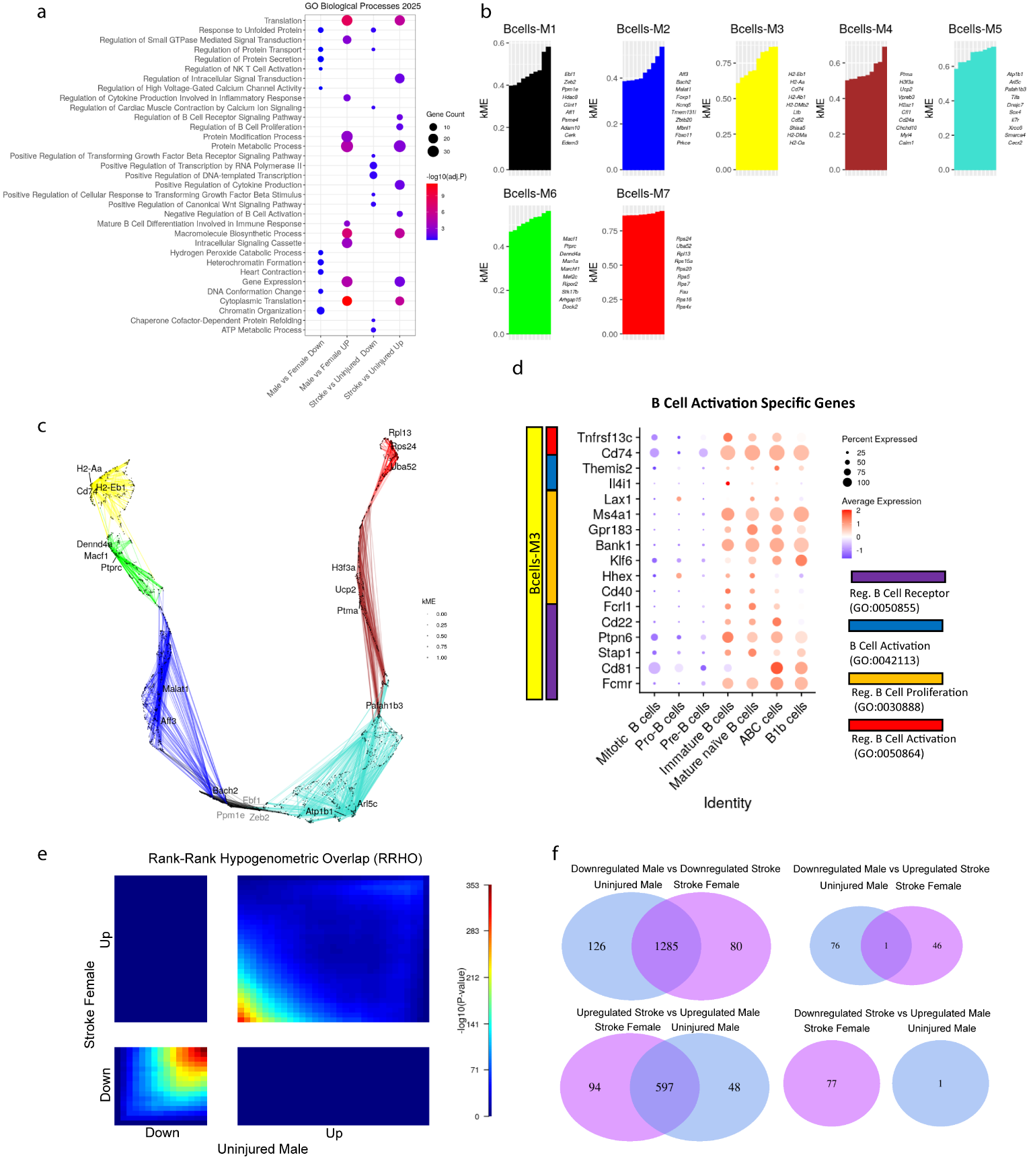

**SFigure 6: B cell GO terms and hdWGCNA module descriptions**

**(a)** Dotplot of gene ontology (GO) Biological Process 2025 terms for uninjured males vs uninjured females and stroke females vs uninjured females **(b)** hdWGCNA 7 Modules (M1 – M7) and their top 10 genes contributing to eigengene based connectivity (kME) per module. **(c)** UMAP of hdWGCNA B cell 7 modules with 3 hub genes labeled for each module. **(d)** Dot plot of B cell genes from module M3 (yellow) relating to B cell activation Go terms (0050855, 0042113,0030888,0050864) across B cell phenotypes show a high expression in the majority of Immature, Mature, and ABC/B1b populations. **(e)** Rank-Rank Hypergeometric Overlap (RRHO) heatmap plot of DEG lists ranked by signed p-values of uninjured males vs uninjured females (Uninjured Males) and stroke females vs uninjured females (Stroke Females). **(f)** RRHO Venn diagrams of all combinations of gene regulations: down-down (top left), up-down (top right), up-up (bottom left), down-up (bottom right) for uninured male (blue) and stroke female (pink).

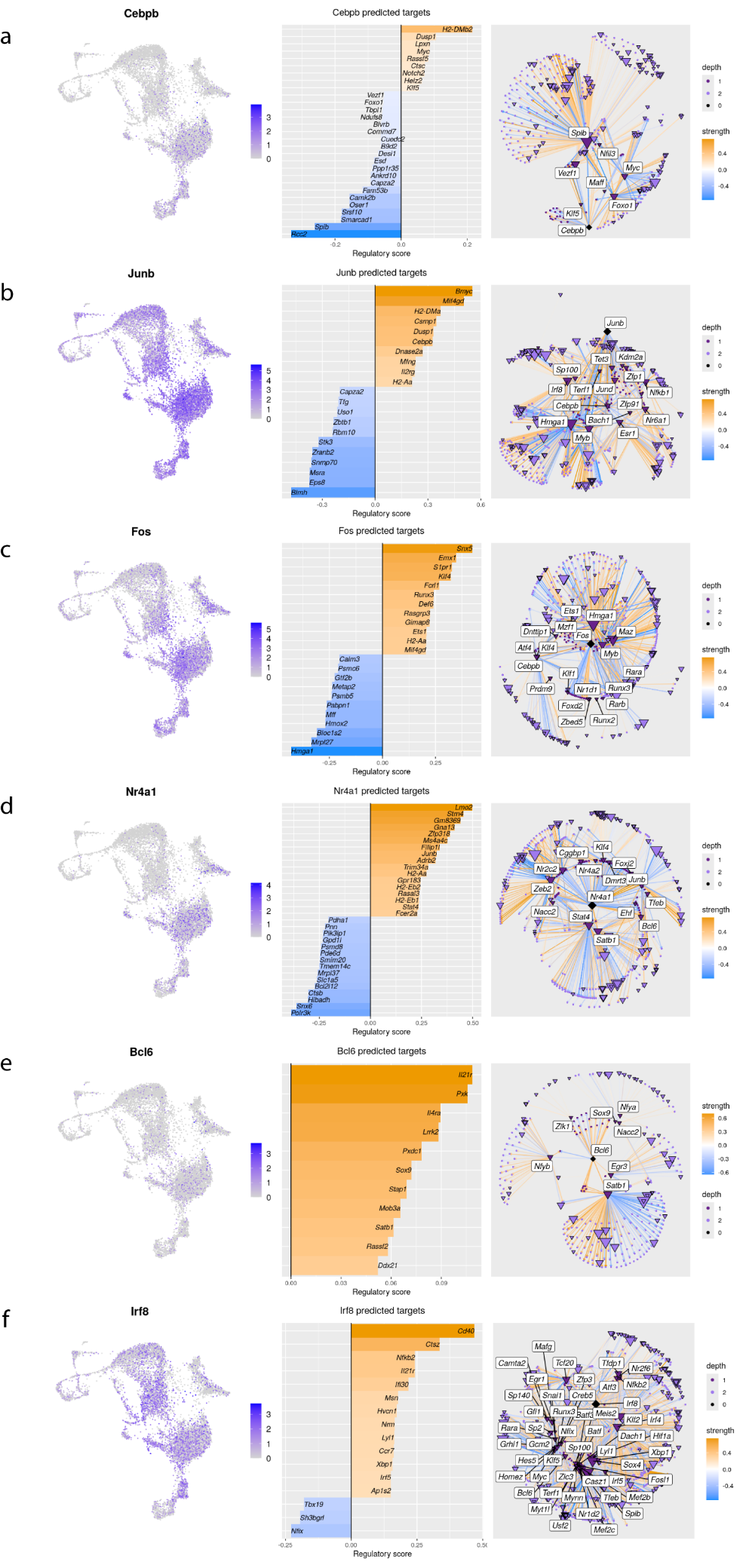

**SFigure 7: Transcription factor regulation networks of significantly regulated TFs**

Transcription Factor (TF) regulon plots for 6 significantly regulated TFs. On the **Left** of each panel is the Gene feature plot superimposed onto all B cell populations, in the **Middle** are the Top regulated genes by TF ranked by regulatory score with orange being genes activated by the TF and blue bars denoting genes suppressed by the TF , while on the **Right** of each panel if the directed TF network plot of the TF with their targets (depth 2) for: **(a)** *Cebpb*; **(b)** *Junb*; **(c)** *Fos*; **(d)** *Nr4a1*; **(e)** *Bcl6*; and **(f)** *Irf8*.

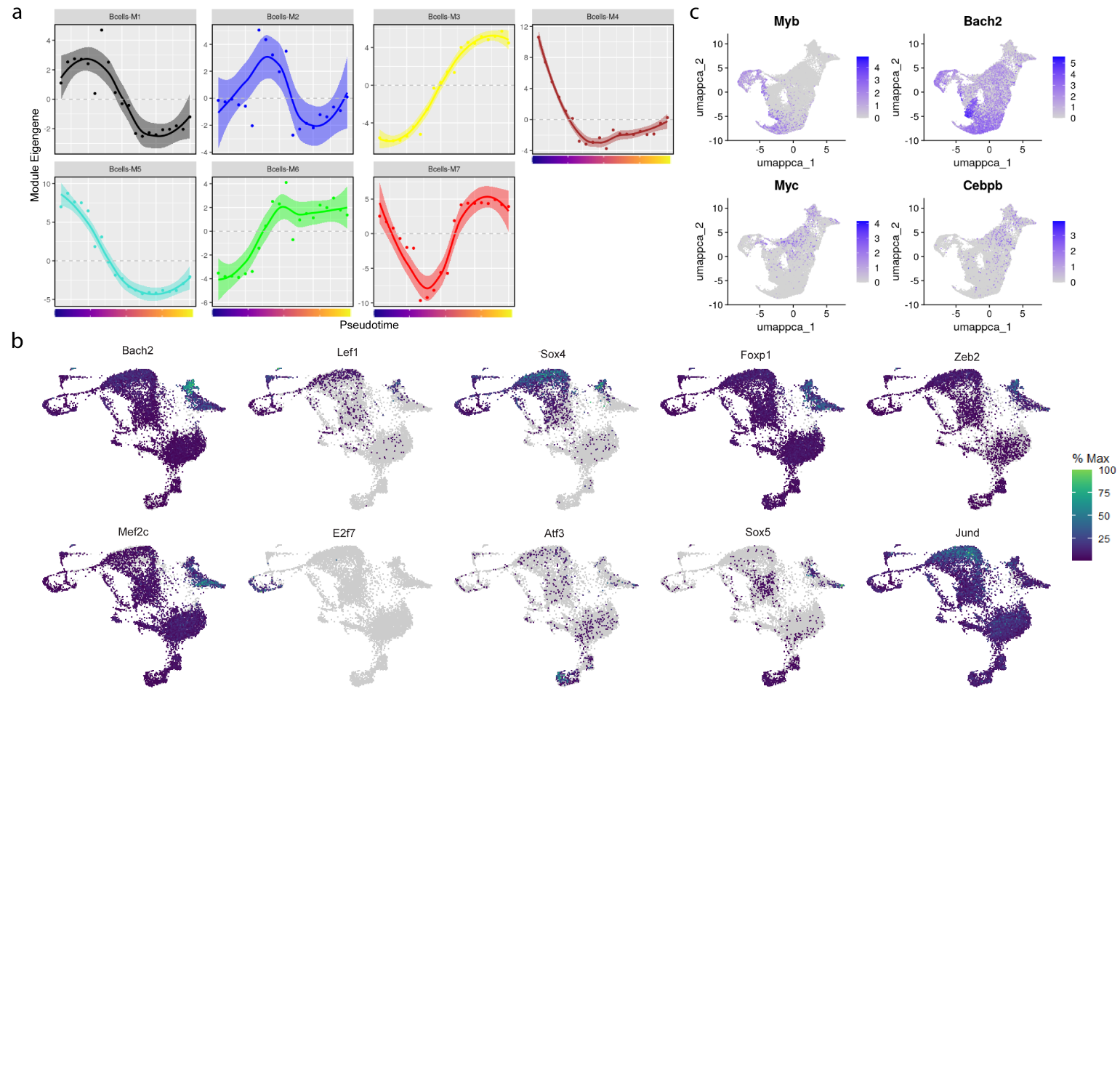

**SFigure 8: Differential pseudotime trajectory analysis of hdWGCNA modules and transcription factors**

**(a)** hdWGCNA B cell module eigengene dynamics plotted across Monocole3 ordered pseudotime. **(b)** Pseudotime differential gene expression analysis via Monocole3 (P value < 0.05) filtering for top 12 transcription factors in order of Moran’s I statistic. **(c)** Gene feature plots of *Myb*, *Bach2*, *Myc*, & *Cebpb* highlight the activation of TF in ABC/B1b populations.

**Supplemental Tables**

**STable 1. Complete 3-way rmANOVA statistics for flow cytometry shown in Figure 1 and SFig. 1**

| **CD45+ Leukocytes (SFig. 1a)** | |  |  |  |
| --- | --- | --- | --- | --- |
| **Source of Variation** | **F (DFn, DFd)** | **% total variation** | **P value** |  |
| Brain Section | F (1.320, 39.59) = 4.739 | 7.75 | 0.0263 | * |
| Sex | F (1, 30) = 5.989 | 6.353 | 0.0205 | * |
| Injury | F (1, 30) = 1.206 | 1.28 | 0.2808 | ns |
| Brain Section x Sex | F (1.320, 39.59) = 0.08093 | 0.1324 | 0.8439 | ns |
| Brain Section x Injury | F (1.320, 39.59) = 1.259 | 2.059 | 0.2814 | ns |
| Sex x Injury | F (1, 30) = 2.338 | 2.48 | 0.1367 | ns |
| Brain Section x Sex x Injury | F (1.320, 39.59) = 0.03412 | 0.0558 | 0.9106 | ns |
| **CD19+ B cells (SFig. 1b)** | |  |  |  |
| **Fixed effects (type III)** | **F (DFn, DFd)** |  | **P value** |  |
| Brain Section | F (1.855, 54.73) = 2.379 |  | 0.1059 | ns |
| Sex | F (1, 30) = 3.696 |  | 0.0641 | ns |
| Injury | F (1, 30) = 4.767 |  | 0.037 | * |
| Brain Section x Sex | F (1.855, 54.73) = 3.358 |  | 0.0455 | * |
| Brain Section x Injury | F (1.855, 54.73) = 0.8339 |  | 0.432 | ns |
| Sex x Injury | F (1, 30) = 0.5744 |  | 0.4544 | ns |
| Brain Section x Sex x Injury | F (1.855, 54.73) = 4.182 |  | 0.0229 | * |
| **CD11b+ B cells (SFig. 1c)** | |  |  |  |
| **Source of Variation** | **F (DFn, DFd)** | **% of total variation** | **P value** |  |
| Brain Section | F (1.691, 50.73) = 4.179 | 5.77 | 0.0265 | * |
| Sex | F (1, 30) = 7.821 | 9.517 | 0.0089 | ** |
| Injury | F (1, 30) = 1.728 | 2.102 | 0.1987 | ns |
| Brain Section x Sex | F (1.691, 50.73) = 0.6182 | 0.8536 | 0.5167 | ns |
| Brain Section x Injury | F (1.691, 50.73) = 0.9961 | 1.375 | 0.3646 | ns |
| Sex x Injury | F (1, 30) = 1.170 | 1.423 | 0.2881 | ns |
| Brain Section x Sex x Injury | F (1.691, 50.73) = 1.017 | 1.404 | 0.3578 | ns |
| **CD11c+ B cells (Fig. 1a)** |  |  |  |  |
| **Source of Variation** | **F (DFn, DFd)** | **% of total variation** | **P value** |  |
| Brain Section | F (1.713, 51.38) = 6.263 | 5.851 | 0.0055 | ** |
| Sex | F (1, 30) = 5.463 | 8.406 | 0.0263 | * |
| Injury | F (1, 30) = 3.340 | 5.139 | 0.0776 | ns |
| Brain Section x Sex | F (1.713, 51.38) = 1.923 | 1.797 | 0.1619 | ns |
| Brain Section x Injury | F (1.713, 51.38) = 3.476 | 3.248 | 0.0451 | * |
| Sex x Injury | F (1, 30) = 1.302 | 2.003 | 0.263 | ns |
| Brain Section x Sex x Injury | F (1.713, 51.38) = 1.625 | 1.518 | 0.2093 | ns |
| **CD11b+CD11c+ B cells (Fig. 1b)** | |  |  |  |
| **Source of Variation** | **F (DFn, DFd)** | **% of total variation** | **P value** |  |
| Brain Section | F (1.529, 45.86) = 5.288 | 4.671 | 0.0142 | * |
| Sex | F (1, 30) = 3.891 | 5.973 | 0.0578 | ns |
| Injury | F (1, 30) = 5.979 | 9.178 | 0.0206 | * |
| Brain Section x Sex | F (1.529, 45.86) = 2.273 | 2.008 | 0.1262 | ns |
| Brain Section x Injury | F (1.529, 45.86) = 3.971 | 3.507 | 0.0357 | * |
| Sex x Injury | F (1, 30) = 2.501 | 3.839 | 0.1243 | ns |
| Brain Section x Sex x Injury | F (1.529, 45.86) = 1.516 | 1.339 | 0.2308 | ns |
| **CD23+CCR7+ B cells (Fig. 1c)** | |  |  |  |
| **Fixed effects (type III)** | **F (DFn, DFd)** |  | **P value** |  |
| Brain Section | F (1.089, 32.13) = 5.749 |  | 0.0202 | * |
| Sex | F (1, 30) = 4.524 |  | 0.0418 | * |
| Injury | F (1, 30) = 4.513 |  | 0.042 | * |
| Brain Section x Sex | F (1.089, 32.13) = 6.338 |  | 0.015 | * |
| Brain Section x Injury | F (1.089, 32.13) = 4.943 |  | 0.0306 | * |
| Sex x Injury | F (1, 30) = 5.991 |  | 0.0204 | * |
| Brain Section x Sex x Injury | F (1.089, 32.13) = 6.277 |  | 0.0155 | * |
| **Plasma cells (Fig. 1d)** |  |  |  |  |
| **Fixed effects (type III)** | **F (DFn, DFd)** |  | **P value** |  |
| Brain Section | F (1.189, 35.08) = 4.268 |  | 0.04 | * |
| Sex | F (1, 30) = 1.779 |  | 0.1923 | ns |
| Injury | F (1, 30) = 6.589 |  | 0.0155 | * |
| Brain Section x Sex | F (1.189, 35.08) = 6.410 |  | 0.0122 | * |
| Brain Section x Injury | F (1.189, 35.08) = 3.049 |  | 0.0832 | ns |
| Sex x Injury | F (1, 30) = 2.181 |  | 0.1502 | ns |
| Brain Section x Sex x Injury | F (1.189, 35.08) = 7.200 |  | 0.0081 | ** |
| **T cells (SFig. 1d)** | |  |  |  |
| **Source of Variation** | **F (DFn, DFd)** | **% of total variation** | **P value** |  |
| Brain Section | F (1.680, 50.40) = 7.275 | 7.206 | 0.0028 | ** |
| Sex | F (1, 30) = 1.734 | 3.131 | 0.1979 | ns |
| Injury | F (1, 30) = 0.4710 | 0.8506 | 0.4978 | ns |
| Brain Section x Sex | F (1.680, 50.40) = 1.489 | 1.475 | 0.2356 | ns |
| Brain Section x Injury | F (1.680, 50.40) = 1.493 | 1.479 | 0.2348 | ns |
| Sex x Injury | F (1, 30) = 0.4075 | 0.7359 | 0.5281 | ns |
| Brain Section x Sex x Injury | F (1.680, 50.40) = 0.6126 | 0.6067 | 0.5185 | ns |

**STable 2**. **Statistics for behavioral comparisons with flow cytometry populations in Figure 1**

| **3-way rmANOVA for Open Field - Time in Periphery (%) (Extended Data Fig. 1e)** | | | | |
| --- | --- | --- | --- | --- |
| **Fixed effects (type III)** | **F (DFn, DFd)** |  | **P value** |  |
| Day | F (2.640, 73.03) = 2.874 |  | 0.0484 | * |
| Sex | F (1, 28) = 1.403 |  | 0.2461 | ns |
| Injury | F (1, 28) = 2.453 |  | 0.1286 | ns |
| Day  x Sex | F (2.640, 73.03) = 0.4986 |  | 0.6608 | ns |
| Day  x Injury | F (2.640, 73.03) = 1.656 |  | 0.1893 | ns |
| Sex x Injury | F (1, 28) = 3.345 |  | 0.0781 | ns |
| Day  x Sex x Injury | F (2.640, 73.03) = 1.184 |  | 0.3192 | ns |
| **3-way rmANOVA for Open Field - Distance Travelled (Extended Data Fig. 1f)** | | | | |
| **ANOVA table** | **F (DFn, DFd)** | **% of total variation** | **P value** |  |
| Day | F (1.822, 51.03) = 13.79 | 15.56 | <0.0001 | **** |
| Sex | F (1, 28) = 3.199 | 4.164 | 0.0845 | ns |
| Injury | F (1, 28) = 4.906 | 6.385 | 0.0351 | * |
| Day  x Sex | F (1.822, 51.03) = 0.9978 | 1.126 | 0.3693 | ns |
| Day  x Injury | F (1.822, 51.03) = 2.629 | 2.967 | 0.0866 | ns |
| Sex x Injury | F (1, 28) = 1.445 | 1.881 | 0.2393 | ns |
| Day  x Sex x Injury | F (1.822, 51.03) = 2.678 | 3.022 | 0.0831 | ns |
| **3-way rmANOVA 10 day Baseline for Rotarod (RR) (Extended Data Fig. 1g)** | | | | |
| **ANOVA table** | **F (DFn, DFd)** | **% of total variation** | **P value** |  |
| Day | F (2.921, 81.80) = 17.90 | 9.329 | <0.0001 | **** |
| Sex | F (1, 28) = 0.6054 | 1.476 | 0.4431 | ns |
| Injury | F (1, 28) = 1.306 | 3.183 | 0.2628 | ns |
| Day x Sex | F (2.921, 81.80) = 0.3700 | 0.1929 | 0.7696 | ns |
| Day x Injury | F (2.921, 81.80) = 0.3932 | 0.2049 | 0.753 | ns |
| Sex x Injury | F (1, 28) = 0.8041 | 1.96 | 0.3775 | ns |
| Day x Sex x Injury | F (2.921, 81.80) = 0.4749 | 0.2475 | 0.6956 | ns |
| **3-way rmANOVA Baseline vs. post-stroke for Rotarod (Fig. 1f)** | | |  |  |
| **Fixed effects (type III)** | **F (DFn, DFd)** |  | **P value** |  |
| Day | F (2.016, 55.09) = 4.638 |  | 0.0136 | * |
| Sex | F (1, 28) = 0.2253 |  | 0.6387 | ns |
| Injury | F (1, 28) = 4.149 |  | 0.0512 | ns |
| Day  x Sex | F (2.016, 55.09) = 1.133 |  | 0.3298 | ns |
| Day  x Injury | F (2.016, 55.09) = 3.528 |  | 0.0359 | * |
| Sex x Injury | F (1, 28) = 2.359 |  | 0.1358 | ns |
| Day  x Sex x Injury | F (2.016, 55.09) = 1.187 |  | 0.313 | ns |
| **RR at 1 wk vs. plasma cells at 3 wks post-stroke (Fig. 1g top row)** | | | |  |
|  | **Females** | **Males** |  |  |
| F | 2.831 | 7.779 |  |  |
| DFn, DFd | 1, 8 | 1, 4 |  |  |
| P value | 0.131 | 0.0494 |  |  |
| Deviation from zero? | Not Significant | Significant |  |  |
| **RR at 3 wks vs. plasma cells at 3 wks post-stroke (Fig. 1g bottom row)** | | | |  |
|  | **Females** | **Males** |  |  |
| F | 9.215 | 136.3 |  |  |
| DFn, DFd | 1, 8 | 1, 3 |  |  |
| P value | 0.0162 | 0.0013 |  |  |
| Deviation from zero? | Significant | Significant |  |  |

**STable 3**. **Statistics for progenitor B cell populations in the cerebellum (Fig. 2f)**

| **Pre Pro B cells** |  |  |  |
| --- | --- | --- | --- |
| Source of Variation | % of total variation | P value |  |
| Sex | 19.23 | 0.0518 | ns |
| Injury | 27.3 | 0.0235 | * |
| **Pro B cells** |  |  |  |
| Source of Variation | % of total variation | P value |  |
| Sex | 2.288 | 0.4216 | ns |
| Injury | 21.15 | 0.0233 | * |
| **Pre B cells** |  |  |  |
| Source of Variation | % of total variation | P value |  |
| Sex | 10.28 | 0.1385 | ns |
| Injury | 4.203 | 0.3336 | ns |
| **Immature B cells** |  |  |  |
| Source of Variation | % of total variation | P value |  |
| Sex | 4.065 | 0.3757 | ns |
| Injury | 5.346 | 0.3116 | ns |

**STable 4. Clinical and demographic characteristics of study donors** **from the rapid postmortem brain donation program at the University of Kentucky (Fig. 6)**

| **ID** | **Age** | **Sex** | **Cognitive status** | **APOE** | **History of Stroke** |
| --- | --- | --- | --- | --- | --- |
| 1 | 96 | Female | MCI | ε3/ε3 | No |
| 2 | 68 | Female | Demented | N/A | No |
| 3 | 89 | Male | Demented | ε3/ε3 | No |
| 4 | 83 | Female | Normal | ε3/ε3 | Yes |
| 5 | 91 | Male | Demented | ε3/ε3 | No |
| 6 | 84 | Male | Normal | ε3/ε4 | No |
| 7 | 100 | Female | Demented | ε3/ε3 | No |
| 8 | 81 | Male | Demented | ε3/ε4 | No |
| 9 | 83 | Female | Demented | ε3/ε4 | No |
| 10 | 90 | Female | Normal | ε2/ε3 | No |
| 11 | 82 | Male | Demented | ε3/ε3 | Yes |
| 12 | 86 | Male | Demented | ε3/ε3 | No |
| 13 | 82 | Male | Demented | ε3/ε4 | Yes |
| 14 | 84 | Male | MCI | ε3/ε4 | No |

Cognitive status was defined in a range of normal, mild-cognitive impairment (MCI) or Demented; APOE, apolipoprotein E.

**STable 5. Statistical interactions for human parenchymal ABC data by sex and age (Figure 6)**

| **2way rmANOVA of CD11b+ ABCs (Fig. 6c)** | | |  |  |
| --- | --- | --- | --- | --- |
| **ANOVA table** | **F (DFn, DFd)** | **% of total variation** | **P value** |  |
| Section x Sex | F (1, 11) = 1.086 | 3.069 | 0.3197 | ns |
| Section | F (1, 11) = 0.2448 | 0.6917 | 0.6305 | ns |
| Sex | F (1, 11) = 3.753 | 16.63 | 0.0788 | ns |
| Subject | F (11, 11) = 1.568 | 48.73 | 0.234 | ns |
| **2way rmANOVA of CD11c+ ABCs (Fig. 6d)** | | |  |  |
| **ANOVA table** | **F (DFn, DFd)** | **% of total variation** | **P value** |  |
| Section x Sex | F (1, 11) = 2.529 | 3.981 | 0.1401 | ns |
| Section | F (1, 11) = 0.7838 | 1.234 | 0.3949 | ns |
| Sex | F (1, 11) = 0.07564 | 0.5265 | 0.7884 | ns |
| Subject | F (11, 11) = 4.421 | 76.57 | 0.0104 | * |
| **2way rmANOVA of CD23+ Mature B cells (Fig. 6e)** | | |  |  |
| **ANOVA table** | **F (DFn, DFd)** | **% of total variation** | **P value** |  |
| Section x Sex | F (1, 11) = 0.6793 | 1.363 | 0.4273 | ns |
| Section | F (1, 11) = 0.5512 | 1.106 | 0.4734 | ns |
| Sex | F (1, 11) = 0.001797 | 0.01236 | 0.9669 | ns |
| Subject | F (11, 11) = 3.428 | 75.63 | 0.0262 | * |
| **CD11b+ ABCs in the cerebellum vs. Age (Fig. 6f)** | | |  |  |
|  | **Females** | **Males** |  |  |
| F | 0.2619 | 0.000003181 |  |  |
| DFn, DFd | 1, 4 | 1, 5 |  |  |
| P value | 0.6358 | 0.9986 |  |  |
| Deviation from zero? | Not Significant | Not Significant |  |  |
| **CD11c+ ABCs in the cerebellum vs. Age (Fig. 6g)** | | |  |  |
|  | **Females** | **Males** |  |  |
| F | 0.5957 | 9.157 |  |  |
| DFn, DFd | 1, 4 | 1, 5 |  |  |
| P value | 0.4833 | 0.0292 |  |  |
| Deviation from zero? | Not Significant | Significant |  |  |
